# Uncovering the domain language of protein function and protein networks using DANSy

**DOI:** 10.1101/2024.12.04.626803

**Authors:** Adrian A. Shimpi, Kristen M. Naegle

## Abstract

Protein-protein interaction networks can help identify co-regulated modules, emergent biology (such as pathways), disease-associated partners, and function through association. However, these networks are limited by the breadth of experimental data behind them, which is incomplete and uneven across the proteome. For example, the coverage of interactions driven by reversible post-translational modifications (acetylation, phosphorylation, etc.) or protein fusions in diseases are particularly difficult to establish. Protein domains, conserved structural and functional units, are a major component of what defines a proteins function and its interactions. Whereas protein interactions are sparsely understood at this time, domain identification and coverage is mature. In this work, we propose a language-based network that utilizes the domain as “words” that makes up protein “sentences” to cover an entire proteome. We first convert proteins into n-grams (a formalization of contiguous words in a sentence) and assemble n-grams into a comprehensive network. We then use information theory to reduce the complexity of the network, collapsing to a network and n-gram size that recovers the majority of the proteome complexity. Using network theoretic approaches, we explore the larger human proteome and subnetwork analysis to understand specific properties in two applications: reversible systems of post-translational modifications (PTMs) and cancer fusion genes. PTM analysis across species suggests that reversible PTM systems convergently evolved similar domain architectures – allowing higher interconnectivity between reader and writer domains, while eraser domains remained highly disconnected. Cancer fusion analysis finds that, despite the possibility that fusions may sample novel domain word combinations, creating new connections or altering the human protein network, most cancer fusion genes follow existing domain combination rules. Collectively, these results suggest that an n-gram based analysis of proteomes complements direct protein interaction approaches, but provides a more fully described network of interconnected protein function that can provide unique insights on signaling pathway analysis. We refer to this approach for converting proteomes into functional linguistic networks as Domain Architecture Network Syntax (DANSy).

**Significance:** Here, we develop a novel computational method that treats proteins as sentences made up of domain words. We integrate this linguistic representation with networks to uncover abstract functional relationships to more fully represent the proteome than traditional protein-protein interaction networks, which are limited by incomplete experimental data. Our findings show how this framework, which we term Domain Architecture Network Syntax (DANSy), uncovers common “grammar” governing protein functionality in the evolution of post-translational modification systems and demonstrates how cancer fusion genes maintain established principles of the natural proteome.

## Introduction

Domains are modular units of structure and function that enable protein complex formation or translate biochemical information between signaling effectors [1]. About half of the human proteome consists of multi-domain proteins, where domains help define overall protein functionality through their independent contributions. The combination of domains within a protein, or domain architecture, arises primarily through the shuffling of pre-existing domains, rather than the emergence of new domains [2–4]. Changes in domain architectures most commonly occur from the gain or loss of domains at the terminal ends of pre-existing proteins [5]. The domain architectures of protein families encode evolutionary jumps and can predict the acquisition of new protein functions [3, 6]. Interestingly, only a small fraction of possible domain combinations are observed, which cannot be described by random shuffling of domains during genetic recombination events [3, 4, 7]. Instead, the restrictions on domain combinations are likely attributed to few domains having multiple feasible domain partners [8, 9]. These observations have motivated representing proteins as vectors of their domains to evaluate the evolution of protein families and the complete proteome [3, 6, 10]. These representations can predict the subcellular localization and gene ontology terms for individual proteins [11, 12], suggesting protein domain architectures encode several layers of functional and evolutionary information.

Insights from broad surveys of both protein domain architectures and amino acid sequences suggest that proteins operate with sets of rules akin to grammar within natural languages [10, 13, 14]. Protein language models often focus on the amino acid sequence to predict the structure, function, and evolution of unresolved proteins or guide novel protein design [13, 15–17]. Despite the breadth of information encoded within domain architectures few linguistic approaches have used domains as the fundamental unit of the protein language. Applications that utilize domains as the fundamental units of analysis can recover protein functionality and evolution across the tree of life, independent of known protein interaction networks or signaling pathways [6, 10]. However, these past methods primarily focus on the occurrence of a domain in a protein rather than its sequential ordering and repetition. The sequential order of domains likely reflect the evolutionary pressures that encourage the non-random shuffling of domains. This suggests the relative location and combination of certain domain combinations constrain and determine protein function. For example, catalytic domains, like kinase domains, can occur on their own, but can also be found in domain architectures with binding domains (e.g. SH2 or WW domains), which changes its specificity by creating avidity effects [18, 19]. However, certain types of activity-modifying domains, or combinations of catalytic domains, may be prohibited to ensure feasibility and interpretability within the biochemical networks of the cell, akin to sentence clarity within natural languages. In this work, we were interested in learning possible global constraints on such domain syntax rules in the human proteome and their implications for specific functional systems.

N-gram analysis is a linguistic approach that maintains both the composition and sequential order of words within natural languages and can be readily adapted to protein domain architectures. By treating individual domains as words and the complete domain architecture of a protein as a sentence, n-grams with n domains can be extracted from either single or multiple domain architectures. A 2-gram model has shown that pairwise domain combinations can recapitulate the evolution of proteome complexity [10]. The smaller fraction of multidomain proteins in prokaryotes than eukaryotes [20] has limited n-gram analysis to mostly 2-gram models for comparative genomic studies. However, certain 2-3 domain combinations, or supra-domains, are rearranged together as a unit across protein families [21]. If an n-gram model does not extract n-grams longer than these supra-domains, the diversity of feasible domain combinations in a proteome may not be sufficiently recovered. Currently, it is unclear how large n-grams must be to describe the full complexity of the human proteome.

Here, we explore how n-gram models describe the human proteome, independent of sparse protein-protein interactions or related annotations. We combine these models with network analysis techniques to develop a global analysis framework to identify protein domains and multi-domain n-grams that connect obligate domain families for both the proteome and subnetworks within it. We have termed this n-gram network framework as Domain Architecture Network Syntax (DANSy). Importantly, the DANSy network representation relies only on domain annotations, which are more fully annotated than protein-protein interactions [22, 23]. As a result, the connections within the network reflect the related functional connections amongst proteins more than direct protein-protein interactions. We applied DANSy and characterized the entropic information encoded by different domain n-gram lengths needed to recover the complexity of the entire human proteome. Next, we measured the emergent properties of domain combinations in reader-write-eraser systems that coordinate reversible post-translational modifications (PTMs). Surprisingly, we find that despite vastly different evolutionary timescales, most reversible PTM-systems converge to have tight connections between the readers and writers and very loosely, if at all connected, erasers. Looking across evolution, it appears that how PTM systems sampled different word ordering gives a possible measure of the relative time since the appearance of the system and the time at the PTM system rules and composition becomes fixed. We then ask if DANSy analysis can delineate how cancer gene fusions alter, or not, the functional connections within the proteome. Interestingly, we find that predominantly somatic fusion genes arise with the same characteristics that appear to guide the overall evolution of most multidomain protein architectures [3, 4]. Hence, we find that DANSy provides unique insights about the functional connections between proteins and systems within cells at both the whole proteome or in more specialized subsystem networks.

## Results

### Generating domain n-gram networks to describe the proteome

To harness domains to generate a proteome-level functional network using a language-based approach, we first set out to convert proteins into n-grams, which encode contiguous domain-word sequences in all proteins across a proteome. We used UniProt [24] and InterPro [25], via our python-based CoDIAC [26] package, to create a resource of unique canonical proteins and their domains. Each protein is converted into all possible domain n-grams spanning the protein. For example, a three domain protein is converted into six unique n-grams (three 1-grams, two 2-grams, and 1 3-gram). We applied this to the human proteome and analyzed n-gram distributions. Given titin (TTN) is a significant outlier (at 303 domains), we set the maximum number of domains for n-gram extraction to 66, the second-largest number of domains on a human protein, although most of the proteome (95%) has 10 or fewer domains (Fig. 1B, S1A). After encoding, the human proteome (20,420 unique proteins) gave rise to 49,308 unique n-grams. Although some domains have duplicated and recombined with diverse domains, our analysis shows most domain n-grams are unique to single proteins (Fig. 1C), suggesting these are specialized domains or domain sets not shared across proteins with varying biological functions. In contrast, some domains are found in many proteins and many n-grams. For example, protein kinase domains, immunoglobulin subtype 1 (Ig_sub), and zinc finger C2H2 type (Znf-C2H2) are both abundant (high 1-gram count numbers across the proteome) and seen in multiple, higher order n-grams (i.e. partnered in combinations with a diversity of domain partners). This simple analysis highlights that n-gram extraction of proteomes captures the breadth of domain combinations, recombinations, and the connection across functional space, suggesting these n-grams are a powerful way to encode proteome-wide information.

**Figure 1.**
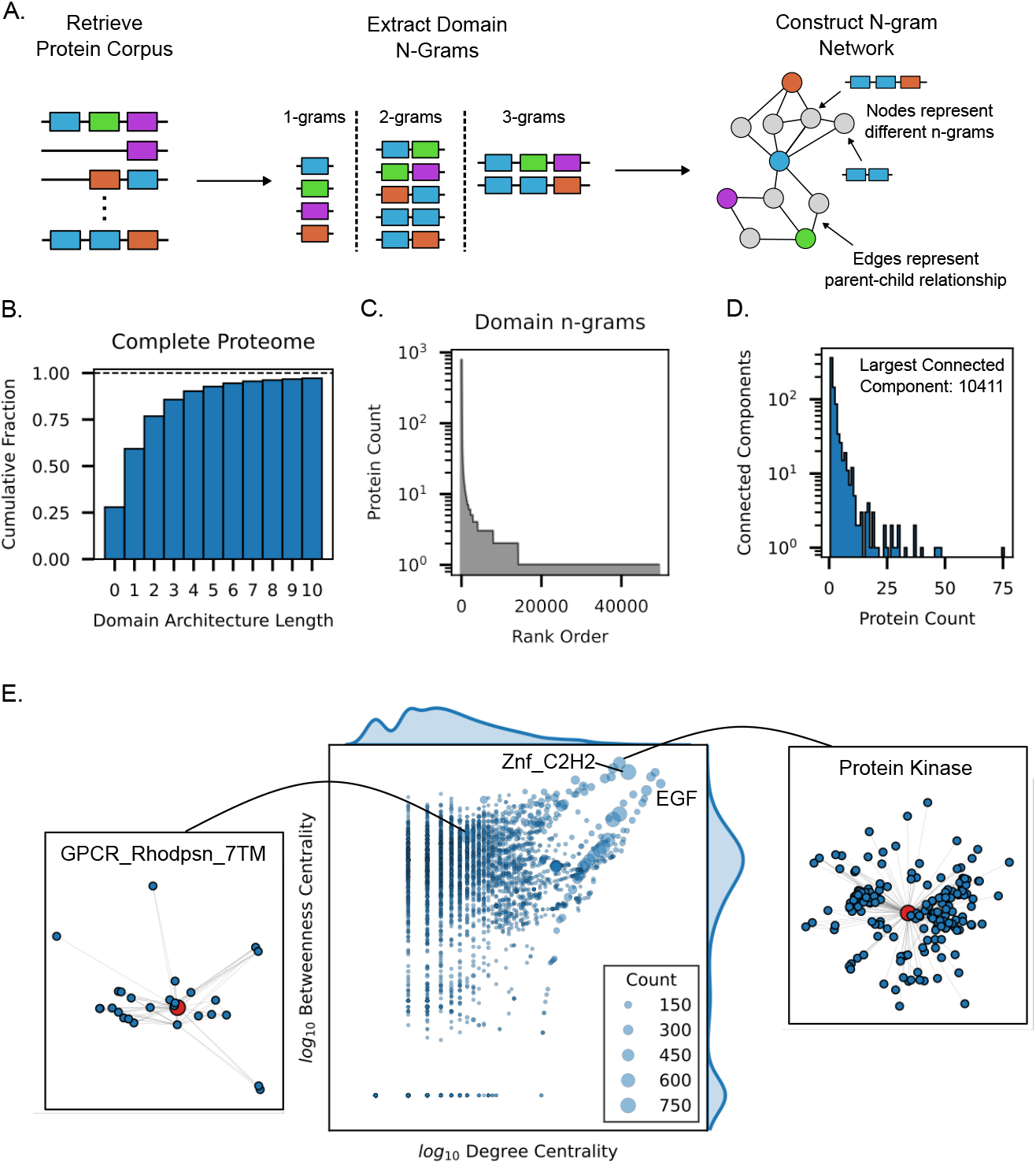
Human proteome n-gram network characterization: A) Overview of DANSy where n-grams of length n are extracted from a collection of protein domain architectures. These n-grams are then used to construct an n-gram network where individual n-grams are nodes and edges represent parent-child relationships where a shorter n-gram is found in a longer n-gram. B) The cumulative distribution of domain architecture lengths across all proteins in the human proteome. C) The number of proteins for each domain n-grams. D) The number of proteins represented for each connected component except the largest connected component which has 10,411 proteins. E) The relationship of each centrality measurement with each other and marker size representing the protein count of each n-gram. For the protein kinase domain and the GPCR_Rhodpsn_7TM domain the subnetworks are displayed with each domain represented by a red node.

We next wished to assemble the entire proteome n-gram language into a single network, an approach we refer to as Domain Architecture Network Syntax (DANSy). Within the DANSy networks, nodes are individual domain n-grams and edges connect nodes with shared n-grams (Fig. 1A). For example, the SH2|Kinase n-gram node has edges to smaller, inclusive 1-grams (SH2 and Kinase 1-gram nodes) and also to a larger architecture, such as the SH3|SH2|Kinase 3-gram node. We constructed a single network of the 49,308 n-grams of the human proteome, allowing us to next use network-based measures to evaluate the human proteome. We proposed that connected components (a gross topographical network feature where a set of nodes only have paths between them and not to any other node in the network) identifies n-gram families that represent domains with shared, compatible properties, since you can traverse amongst these domains through proteins that contain their combinations. The DANSy network of the proteome contained 1494 connected components. Of these connected components, there were 733 isolated nodes (or isolates), which represent singular domains found only within a single family of proteins (e.g. the TACC C-terminal domain within only the TACC1/2/3 proteins). Most non-isolate connected components, which contain 2 or more domain n-grams, represented 5 or fewer proteins except the largest connected component, which represented 10,411 proteins (Fig. 1D). Next, we integrated a node collapsing step during n-gram network construction to reduce redundant n-grams representing the same set of proteins (Fig. S2A), since relatively few domains have a diverse set of domain partners [8] and most domain n-grams represent a single protein or potentially whole protein families. The collapsed network increased the number of isolates to 1219 nodes highlighting an additional 486 unique domain n-grams of specialized functions, which included protein families containing 14 domains (Fig. S2). By creating a non-redundant DANSy n-gram network, we have a foundational representation of the proteome that can be studied to understand general principles or the “grammar” of compatible domain functions and properties.

With the proteome represented by a non-redundant n-gram network, we next set out to use node-centric metrics to identify and understand how specific domains contribute to functional diversity in the proteome. For this, we calculate two complementary centrality metrics: 1) the degree centrality – to identify the domains or domain n-grams with a diverse set of domain partners, and 2) the betweenness centrality – to identify domain n-grams that connect n-gram families by more frequently lying on the shortest paths between nodes in the network. Since 51.0% of the proteome is represented by the largest connected component in the n-gram network, which contains 70.4% of all domain n-grams, we calculated the network metrics only for this set of domain n-grams. Of these analyzed domain n-grams, <1% were within the top 5% of both centrality measurements (Fig. 1E) – this relationship was not strongly associated with the number of proteins containing the domain n-grams. When we compare domain n-grams with different relationships between the centrality measurements we can identify how a domain n-gram contributes to the diversity of protein functionality in the proteome. First, the n-grams with high degree and betweenness centrality values have a diverse set of n-gram partners and connect several n-gram families. These domain n-grams can maintain their functional clarity, but are flexible in which protein contexts they can occur in. Examples of these types of n-grams are the protein kinase, EGF-like, and Znf-C2H2 domains. Second, n-grams with high degree centrality values may have several n-gram partners but frequently have connections within a distinct n-gram family, but rely on another domain n-gram to connect them to the rest of the n-gram network. These n-grams are mostly highly repetitive domain n-grams (e.g. EGF-like repeats with 4 or more copies), which may be found in several proteins, but the biological processes they contribute to are largely driven by their smaller, constituent n-grams. Third, if an n-gram has a high betweenness centrality but low degree centrality it acts as a hub for a small fraction n-grams, which means it is highly constrained in what are compatible domain properties that can modify its general function. An interesting example of this category is the the seven transmembrane region of rhodopsin-like G-protein coupled receptors (GPCR_Rhodpsn_7TM) domain, which is one of the most abundant domains in the proteome but has few feasible domain combinations. If we compare the subnetwork surrounding the GPCR_Rhodpsn_7TM and the protein kinase domain, which is found in a similar number of protein (Fig. S1C), we see how restricted the GPCR_Rhodpsn_7TM domain is in creating multi-domain n-grams, but acts as the localized hub for other smaller n-gram families. Meanwhile, the protein kinase domain is highly connected, and its immediate neighbors connect to other n-grams beyond the kinase domain (Fig. 1E) suggesting it is deeply embedded as a hub for the entire n-gram network. These results demonstrate the power and kinds of insights enabled by DANSy towards describing the extent of compatible functions that underlie many signaling pathways of the proteome.

We next wished to evaluate how large the n-grams must be during extraction and network construction to capture the diversity of the human network. For this analysis, we explored how limiting the n-gram lengths extracted from the proteome impact both the network topology and the changes to the information value represented in the different sets of domain n-grams. Given about 25% of the proteome has 3 or more domains and whole protein families can be represented by up to 14-grams (Fig. 1, S2), we constructed n-gram networks with different maximum n-gram lengths up to 15-grams, and compared this both to the full n-gram network we had already established, and the information content of individual domain frequencies. We found a 10-gram network retains 85% of the entropic information content of the full n-gram network (Supplementary Note 1). Further, the 10-gram model retained all isolates in the network and only lost nodes within the largest connected component representing longer n-grams. Meanwhile, the 2-gram model recreated about 40% of the information content, but introduced 771 new isolates and retained only half of the other non-isolate connected components. These results suggest that only analyzing 2-grams, which have been widely used in prior studies [3, 8, 10], is likely missing key contexts of protein functionality. Alternatively, as we expanded from 10-to 15-gram models, we observed minimal differences between the models, suggesting the increased complexity of longer n-grams did not yield a reciprocal gain in capturing the information content of the proteome (See Supplementary Note 1 for further details). Thus, we selected a 10-gram model to represent the human proteome to balance model complexity and the information content encoded by protein domain architectures.

### Subnetwork analysis to explore reversible post-translational modification systems

We next wished to understand if targeted subnetwork analysis of DANSy helps identify novel information about specific functions. Since interactions in systems like phosphorylation are especially hard to characterize [27] and given the diversity of n-grams containing kinase domains, we wanted to understand if other underlying domain compatibility rules exist for the overall phosphorylation machinery that enables it to regulate most biological processes. Phosphorylation signaling consists of two major subsystems: the phosphotyrosine (pTyr) and the phosphoserine/threonine (pSer/Thr) systems, and both operate under a reader-writer-eraser paradigm to separate the distinct functional modules of the signaling machinery [28]. Since each system uses different domains and there are only a small subset of dual-specificity kinases and phosphatases, we constructed and evaluated n-gram networks for each system separately (domains and classification in Table S1). To understand how interconnected the reader-writer-eraser modules are within each system, we generated two types of n-gram networks: a System Focused model, where n-grams must contain at least one of the domains of the PTM system, and a Complete model, containing all n-grams in each system. The System Focused networks provide insights on how individual components of each system assemble together to maintain or regulate the system’s functionality. Meanwhile, the Complete networks capture the extent of functional reach beyond the main system components. Using these two networks, we can investigate how each system uses domain combinations to define their contributions across biological processes.

First, we focus on the Complete networks to study the extent of external domains used in each system to connect the main system functions to specialized processes. Within the pSer/Thr system, we found the kinase domain extensively combines with more domains than the rest of the system, and few are shared with other system components. Meanwhile, in the pTyr system, the kinase and eraser (PTP) domain share many external n-grams and are highly interconnected. Further, the reader domains remain well-connected to the rest of the network and share several domain n-grams. This interconnectedness establishes the pTyr system as a complete connected graph (i.e. a single connected component), but the pSer/Thr system has multiple connected components, where some n-grams are completely disconnected from the rest of the pSer/pThr machinery. These disconnected n-grams include the phosphatase (eraser), 14-3-3 (reader), and MH2 (reader) domains (Fig. 2A, S9A), which suggests these domains rarely require the same, if any, external domains to define their protein contexts to modulate system functionality. The overlapping domain n-grams between the pTyr components likely suggest these external domains help define the shared biological processes each domain contributes to and regulates. Together, these highlight some key differences in the compatibility of domains that establish the processes each phosphorylation system can regulate.

**Figure 2.**
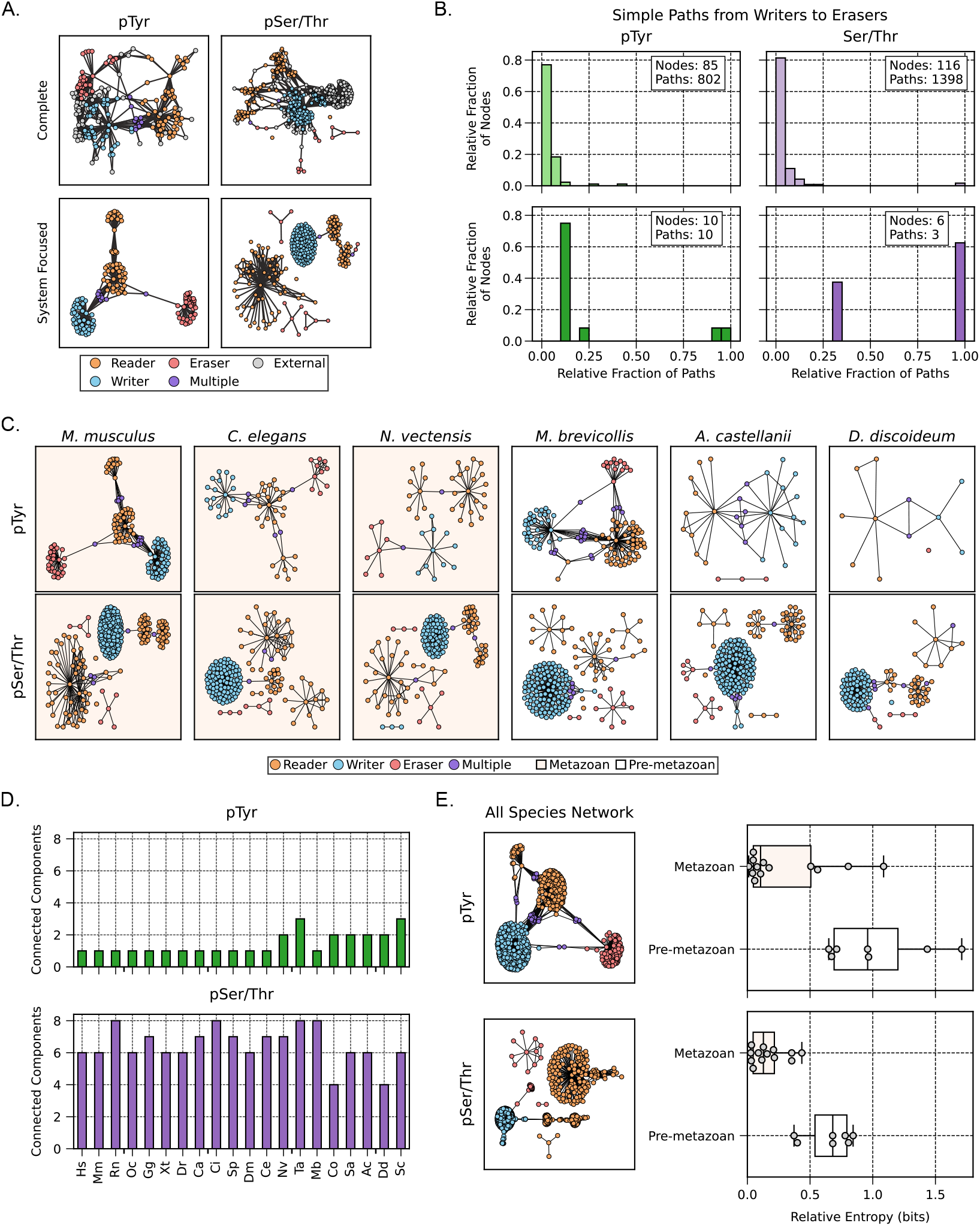
Analyzing the n-gram subnetworks of the pTyr and pSer/Thr systems. A) The n-gram networks for the phosphotyrosine (pTyr) or phosphoserine/threonine (pSer/Thr) systems, where the Complete networks contain all possible n-grams and the System Focused containing only n-grams with the domains of the system. B) The relative fraction of nodes within the simple paths that connect the writer domains to the eraser domains in both phosphorylation systems. C) System Focused n-gram networks of the pTyr or pSer/Thr systems for select species. D) The number of connected components for each phosphorylation system across species. E) The System Focused n-gram network containing all n-grams across all queried species (left) and the relative entropy for metazoans and pre-metazoans relative to the all species network. Hs: *Homo sapiens*, Ms: *Mus musculus*, Rn: *Rattus norvegicus*, Oc: *Oryctolagus cuniculus*, Gg: *Gallus gallus*, Xt: *Xenopus tropicalis*, Dr: *Danio rerio*, Ca: *Carassius auratus*, Sp: *Strongylocentrotus purpuratus*, Dm: *Drosophila melanogaster*, Ce: *Caenorhabditis elegans*, Nv: *Nematostella vectensis*, Ta: *Trichoplax adhaerens*, Mb: *Monosiga brevicollis*, Co: *Capsaspora owczarzaki*, Sa: *Sphaeroforma arctica*, Ac: *Acanthamoeba castellanii*, Dd: *Dictyostelium discoideum*, Sc: *Saccharomyces cerevisiae*

Evaluation of the System Focused networks enable the study of how system components interact and combine to maintain system clarity. Similar to the Complete networks, we see the pTyr network is a complete connected graph, while the pSer/Thr network has multiple connected components. However, unlike the Complete network the pTyr eraser (PTP) domain was only loosely connected to the rest of the network via a single node capturing the SH2|SH2|PTP n-gram of the phosphatases PTPN6 and PTPN11. The SH2 and Tyr Kinase domains remain well-connected, which reflects findings that SH2 domains modulate kinase targeting [19, 29]. Interestingly, in the pSer/Thr system, only one of the previous eraser domains (FCP) remained within the connected component, via the FHA|Kinase and FCP|BRCT domain n-grams. This was the only connected component with a subtype of each reader-writer-eraser module, but unlike the pTyr system there were no shared reader domains between the writer and eraser domains (Fig. 2A,S9A). Given this observation, we next used the simple paths (paths with no repeated nodes) along both the Complete and System Focused networks to further investigate the domains linking the writer domains to eraser domains. For the Complete pSer/Thr network >98% of paths required traversing through the node representing the PH domain, but in the System Focused network all paths required going through 5 of the same nodes (Fig. 2B). Importantly, the paths within the System Focused network can only reach the FCP domain as no other eraser domain shares a common domain partner. In the pTyr networks, there was enough diversity of potential nodes to travel through in the Complete network that no single domain n-gram was found in >45% of all paths (Fig. 2B). However, in the System Focused network all paths to the pTyr eraser domain had to go through the aforementioned SH2|SH2|PTP n-gram. Altogether, these results highlight how external system domains help define the biological context the system can participate in. However, both phosphorylation systems share a common configuration for domain combinations where eraser domains rarely use other system domains to modify their function, while reader domains more readily modulate writer domain activity.

Although we have focused first on analyzing the human proteome, this approach is more generalizable. Therefore, we next wished to explore the evolutionary properties of the pTyr and pSer/pThr signaling systems using DANSy. We built DANSy networks for pTyr and pSer/Thr systems across 20 species, starting from *Saccharomyces cerevisiae*, which contains a single proto-SH2 domain and three PTP domains [30, 31] (Fig. S4). We observe a rapid expansion during the pre-metazoan era in the pTyr n-gram network, but not the pSer/Thr system (Fig. 2C-D,S4,-S7), consistent with what has been observed as the period of rapid expansion for this relatively recently evolved pTyr signaling system [28, 30]. Additionally, we calculated the species-level similarity index of domain n-grams and the relative entropy of the domain n-gram distributions in each species, relative to the collective distribution across all species. Both measures suggest the pSer/Thr networks had already established the stable protein configurations for its tasks by *S. cerevisiae* (Fig. 2E, S8). From the relative entropy measurements, we further observed that most metazoan species had similar distributions as the aggregated species network (designated by lower relative entropy values) for both pTyr and pSer/Thr systems. However, the pSer/Thr system overall had lower relative entropy values. We next wanted to measure properties of system evolution using not only the domain n-gram counts, but the DANSy network configurations. Consistent with rapid evolution period of pTyr signaling, we observed larger configuration differences in pre-metazoan species (Fig. 2D). For example, the pre-metazoan species *Capsaspora owczarzaki* and *Monosiga brevicollis* have several n-grams that connect the PTP domain to the rest of the network, which were lost in metazoans, including n-grams with both kinase and phosphatase domains on single proteins. As metazoans evolved, the SH2|SH2|PTP architecture was the only domain n-gram that persisted, except for *Nematostella vectensis*, one of the earliest metazoans included in our analysis that had additional n-grams connecting the PTP domain to the rest of the system. This suggests that during the rapid expansion of the pTyr system during the emergence of metazoans, the pTyr system was sampling several configurations of domain combinations to create an efficient signaling system.

Since adding the linguistics analysis of domains to the evolutionary analysis of phosphorylation provided new context about how these systems sampled possible combinations to stabilize around specific rules, we next explored additional reversible post-translational modification systems. We generated individual n-gram networks for the acetylation, methylation, and ubiquitination systems, which all follow the reader/writer/eraser paradigm. We constructed a System Focused n-gram network for each system to determine how individual components of each system interact with one another given our results with the phosphorylation systems. Each system had multiple connected components, similar to the pSer/Thr system (Fig. S9). Most of these connected components were single functional modules and frequently involved an eraser domains. If an eraser domain was connected to the rest of the system, it was weakly connected such as the JmjC domain with Tudor domains in the methylation system via the JmjC|Znf_PHD|Znf_PHD|Tudor|Tudor n-gram. These results suggest reversible PTM systems converged to a common set of grammatical rules to maintain clear biochemical functions. Specifically, eraser domains rarely combine with other domains of the PTM system to modify their function.

### Relationship between cancer fusions and protein networks

Having established that network analysis on domain-based n-grams can identify convergent syntax rules in PTM systems, we next explored if DANSy can complement our understanding on how gene fusions possibly act to create novel signaling contexts. Since chimeric proteins produced from gene fusions can have altered functional characteristics than their originating genes [32, 33], we wished to evaluate the domain combinations from predicted chimeric proteins relative to existing combinations in the proteome. We predicted and analyzed chimeric protein domain architectures from >25,000 unique gene fusion breakpoints in 33 study cohorts from the Cancer Genome Atlas (TCGA) within ChimerDB [34]. We mapped the genomic breakpoints to their relative locations in the protein coding sequence and identified the retained domains of each partner gene to build a final predicted domain architecture (Fig. S10). For each predicted domain architecture, we classified fusions if they produced only pre-existing n-grams in the native proteome or created novel domain n-grams. If a fusion gene was predicted to not retain any domains, we labeled these fusions as “no annotation”. Interestingly, fusions that produce novel domain n-grams – to add new nodes and paths in the network – can change the connectivity of the network through multiple mechanisms, but we treat all novel domain architectures as a single category (see Supplementary Note 2). Overall, this collection of chimeric proteins produces a diverse set of domain n-grams that relate the functional properties of gene fusions to impacts on established principles of domain combinations in the proteome.

To characterize the breadth of chimeric protein domain architectures in TCGA, we first analyzed each individual genomic breakpoint for each patient and its network impact on domain linguistics. We stratified the breakpoints and gene fusions into their individual TCGA cohort (i.e. cancer type) and found most breakpoints rarely produced novel domain architectures. Each cohort had less than 20% of breakpoints that result in novel domain architectures except for a few cancer types (thyroid carcinoma THCA, 41.2%, acute myeloid leukemia LAML, 43.8%, and cholangiocarcinoma CHOL, 30.7%). Interestingly, most patient samples had multiple fusion genes and about 39.6% of all patients had multiple breakpoints for at least one fusion gene (Fig. S13A). If a patient sample had multiple breakpoints for the same fusion gene, these largely produced the same domain architecture. Further, less than 5% of samples had fusions with multiple categories (e.g. novel and pre-existing) for the domain architectures (Fig. S13B).

Since most patient samples had chimeric proteins with the same domain architecture even if there were multiple breakpoints, we next analyzed the relationship between the number of breakpoints, unique domain architectures, and unique partner gene pairs in each TCGA cohort. We found in each cohort there were more breakpoints than unique domain architectures and partner gene pairs indicating the presence of recurrent gene fusions and domain architectures (Fig. S12). Since recurrent fusions are a relatively small fraction of gene fusions in the TCGA cohorts [34, 35], we wanted to determine if they more frequently produced novel domain architectures to alter the DANSy networks than singleton fusions. We aggregated gene fusions across the cohorts and classified them on their recurrence across patients, but kept unique domain architectures of the same gene fusion as separate fusions. We found that the recurrent chimeric proteins more readily produce pre-existing domain architectures than fusions that occur in single patients. Further, fusions in the TCGA cohort had fewer novel domain architectures than randomly chosen partner gene pairs and breakpoints (Fig. 3C). These results reinforce that most domains are restricted in what are feasible combinations, and suggests that native domain combinations are favorable for most disease contexts.

**Figure 3.**
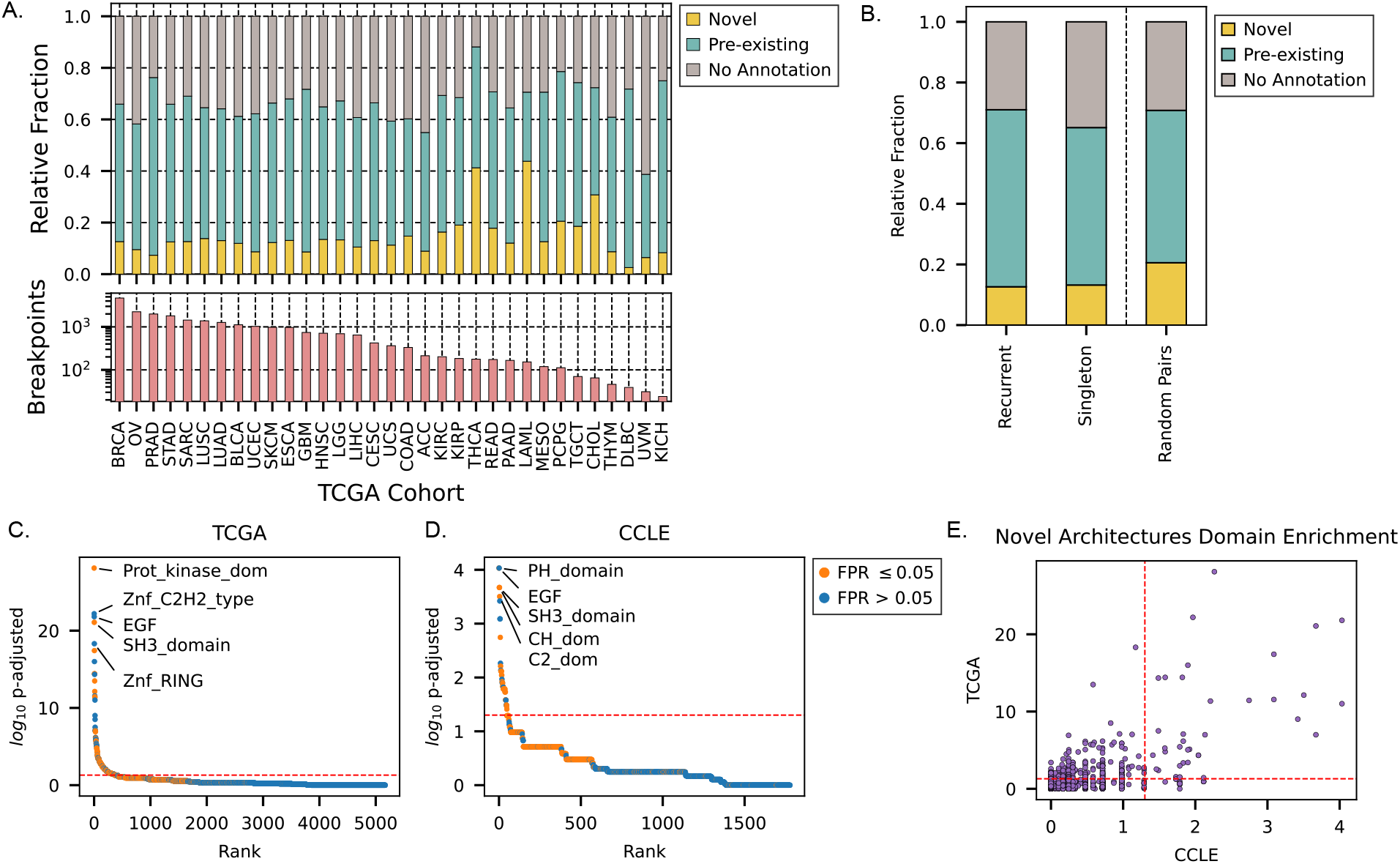
Characterizing predicted protein domain architectures of cancer gene fusions. A) Schematic of the possible changes to the n-gram network caused by individual fusion genes with example fusion genes are predicted domain architectures. For novel domain architectures these can either reinforce existing connected components or reduce the number of connected components. B) The fractional distribution of fusion gene impacts on the n-gram network (top) and the total number of fusion genes (bottom) retrieved for each TCGA study cohort. C) The distribution of architecture n-gram network impacts for recurrent or singleton fusions and a random gene null distribution. D) Domain n-gram enrichment values within novel domain architectures from the TCGA cohort. E) Comparison of domain n-gram enrichment values between the TCGA and CCLE cohorts. Dashed red lines indicate an adjusted p-value of 0.05.

Since fusions with novel domain architectures appear to be an exception, we explored whether they collectively enrich for specific or unique functions (i.e. certain domains). Towards this, we conducted domain enrichment analysis on novel domain architectures and found an enrichment of kinase and DNA binding domains (such as Znf-RING, and Znf-C2H2) in novel fusion genes (Fig. 3D), consistent with prior studies that largely focused on fusion genes in general and not the novel domain architectures [36, 37]. However, several of the top candidates such as the Znf_C2H2 and EGF domains had high false positive rates (≥0.05). We used our centrality metrics to determine if there were associations between a domain’s enrichment in novel domain architectures and its native tendencies to combine or connect different domain n-gram families. Interestingly, several of the most enriched domains were within the top 5% of both centrality measurements within the n-gram network of the entire proteome, but there was no strong correlation between the enrichment values and either centrality value (Fig. S15). This lack of correlation was also observed relative to the number of proteins containing the domain n-gram, suggesting it is not simply how often the domain occurs in the proteome that describes its enrichment in fusion genes. Together, these results highlight the diverse functions chimera proteins can co-opt, irrespective of whether it produces a novel domain architectures. Domains with flexible contexts appear frequently in chimera proteins, but they are not used exclusively to explore new, domain combinations and possibly advantageous functions. These results suggest there is not a preferential protein function being detected across gene fusions, despite the specific classes of gene fusions being frequently identified.

To evaluate if novel domain architectures, or specific domain functions, relate to a fitness advantage, we expanded our analysis to include the Cancer Cell Line Encyclopedia (CCLE), which enables fitness assessment. Similar domain n-grams were enriched in the novel domain architectures across both datasets (Fig. 3C-E), despite only 206 fusion gene-domain architecture pairs being shared between them (Fig. S14A). To relate the domain enrichment and domain architectures to changes in cell fitness levels, we integrated the genome-wide CRISPR loss-of-function screening from the Cancer Dependency Map (DepMap) project to calculate a mean differential dependency scores. For this, we averaged the dependency score of each partner gene for a fusion and calculated the difference in the average score of cell lines with the fusion relative to other cell lines from the same lineage. Most fusions (2082/2143, 97.2%) did not exhibit a dependency for both partner genes (mean dependency score > 0.5), but novel domain architectures showed slightly higher mean dependency scores (0.071 v 0.037, p = 0.10, Fig. S14A). Interestingly, fusions that exhibit dependency had a higher relative fraction of novel domain architectures, but were not enriched for specific domain n-grams (Fig. S14B-D). Together, our results suggest fusion genes not only maintain the established rules of domain combinations of the proteome, but if a new domain combination is created, it reflects a higher dependency on the parent genes, and does not appear to be driven by specific domain functions.

While our DANSy analysis on the DepMap dataset did not suggest specific domains were enriched in fusion genes essential to cancer cell fitness, we found the kinase domain most frequently in cell dependencies, in addition to being highly enriched in novel domain architectures. Kinase fusions have been identified across most cancers [38] with the tyrosine kinases being enriched in gene fusions [36, 39, 40]. Both the TCGA and CCLE datasets had more fusions involving serine/threonine kinases (STKs) genes, however, a larger fraction of the tyrosine kinase (TyrK) fusions resulted in novel domain architectures (Fig. 4A, S16A-B). The kinase domain was retained in ≥ 60% of TyrK fusions and about 25% for STK fusions in both datasets (Fig. 4B, S16C). To understand how different the domain architectures are for both types of kinase fusions, we extracted 1934 domain n-grams from the kinase fusions that retained the kinase domain and found 94.9% of the n-grams were unique to either STK or TyrK fusions including most novel domain n-grams (Fig. 4C, S16E). However, the most prevalent n-grams across all kinase fusions were shared between both STK and TyrK kinase fusions (Fig. 4C, D). These results suggest that while both the TyrK and STK fusions can create novel domain architectures, only a small fraction of domains is shared between the two to explore novel protein functionality.

**Figure 4.**
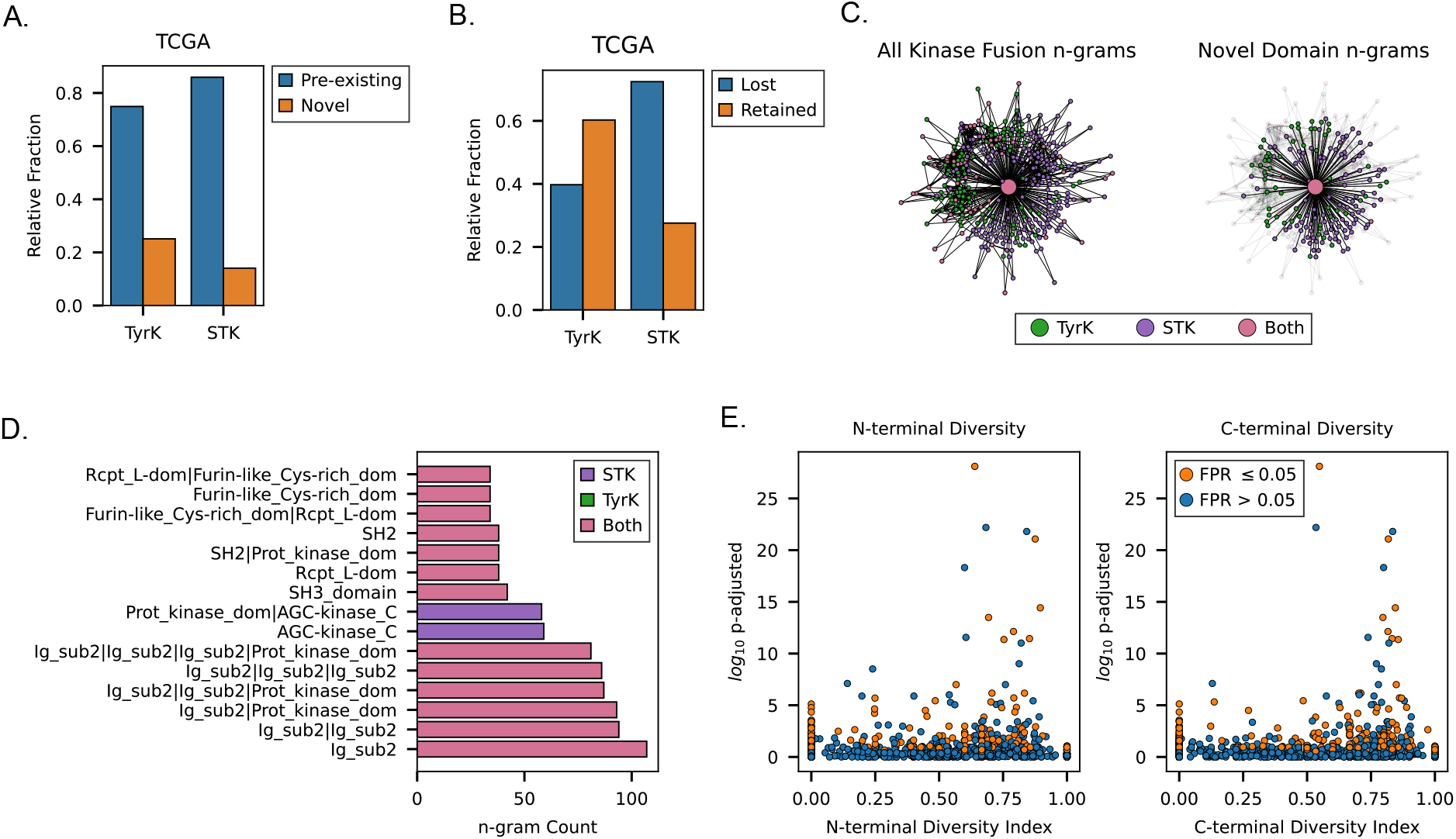
Characterizing predicted protein domain architectures of cancer gene fusions. A) The relative fraction of gene fusions in the TCGA cohort involving kinase domains from either tyrosine kinases (TyrK) or serine/threonine kinases (STK). B) The relative fraction of kinase fusions whose predicted domain architecture retains the protein kinase domain. C) DANSy n-gram network representation of the domain architectures for all kinase fusions that retained a kinase domain with the novel domain combinations shown on the right network. D) The top domain n-grams for kinase fusions. E) The relationship of domain enrichment significance for novel domain architectures relative to the diversity indices for individual domains at either the N-terminal or C-terminal end.

To determine if these observations of novel domain n-grams reflect a natural flexibility of domain partners for each kinase domain, we explored how concentrated natural domain combinations are within the natural proteome (including being the only domain in a protein) for each kinase using the Simpson’s Diversity Index (*D*). We found TyrK domains had fewer domain combinations, but were more evenly distributed (*D* = 0.85) than STK kinase domain (*D* = 0.47) across n-grams towards the N-terminal end. This is largely driven by the large portion of STK (198/402) genes consisting of only a kinase domain. Meanwhile, domains towards the C-terminal side of kinase domains were not extremely diverse (TyrK, *D* = 0.44 and STK, *D* = 0.56) as the kinase domain was frequently the final domain in the domain architecture. Our results suggest the TyrK domain more readily forms novel domain architectures in fusion genes, because it more evenly creates diverse domain n-grams than STK kinase domains, despite there being more STK fusions to increase the chance of forming novel domain n-grams. We asked if the enrichment of a domain in novel domain architectures correlates with the increased diversity of natural domain partners for other domains involved in TCGA fusion genes, but did not find a consistent trend between high enrichment and high diversity indices. However, domains with extremely significant enrichment (p-adjusted ≤ 10^*−*5^) had diversity indices ≥ 0.6 (Fig. 4E) suggesting their inherent flexibility contributes to their ability to form novel domain architectures, but is not the primary determinant.

Collectively, our results demonstrate how DANSy analysis can help uncover aspects of how neofunctionalization of protein domains by gene fusion to rewire signaling pathways in cancer, but also highlights the complexity of gene fusion biology and the need for systemic analyses.

## Discussion

Here, we developed DANSy, which combines domain n-grams with network theory to identify patterns of ordered domain combinations, and explore how these abstract representations of biochemical and structural protein properties connect to each other holistically. Through applications on the human proteome, we identified certain domains, like the protein kinase or Znf-C2H2 domains, act as hubs in the DANSy n-gram network, bridging connections between diverse functional units in the proteome. Network theoretic approaches helped evaluate the complexity needed, in n-gram length, to represent the human proteome, which is significantly larger than prior work focused only on 2-grams. DANSy is versatile and can be applied to different biological contexts to understand how collections of proteins follow sets of principles to coordinate their functionality and establish the system in question. By relying only on protein domain architectures, DANSy can capture functional interactions that are not reliant on annotations that describe known protein-protein interactions or involvement in a biological process. DANSy is another powerful tool for analyzing signaling networks and proteomes to understand the principles guiding protein functionality.

As a demonstration of the insights DANSy can provide, we applied it to understand the rules guiding feasible domain combinations in reversible PTM system. We found that the phosphorylation, methylation, acetylation, and ubiquitination systems converged to a common set of principles where eraser domains rarely utilize other system components to modify their function. When we compared the different phosphorylation systems, we observed that as the pTyr system rapidly expanded, species were sampling several domain combinations suggesting a process of configuring and refining the compatibility rules of protein functionality. This period was near the development of metazoans and was when the separation of catalytic domains was solidified. This suggests that the n-gram network properties can potentially measure if a system is converging to a set of grammatical rules or rapidly testing new domain combinations. Since domains encode the evolutionary jumps of protein families [6], we can use DANSy to study if other biological systems were under selective pressures to explore, diversify, or collapse compatible protein functions to clarify the interactions of different system components. This work presents interesting new questions with regard to reversible PTM systems – e.g. what is the advantage of connecting writer and reader domains, but loosely, or fully disconnecting, eraser domains from readers?

Cancer is considered an evolutionary process where aberrant signaling that increases tumor cell fitness can take on several forms including the generation of fusion genes. However, fusion genes have a complicated role in cancer as they can create signaling dependencies, but most are considered passenger aberrations rather than putative drivers of most cancers [32, 39, 41, 42]. The continued maturation of sequencing technologies and gene fusion detection programs have identified thousands of fusion genes [34, 35, 43], but has created challenges in screening fusion candidates for potential actionable courses of treatment. Here, we aimed to use DANSy to characterize the landscape of domain architectures being generated by fusion genes to attempt to prioritize candidates, but surprisingly found most predicted fusion gene domain architectures recreated pre-existing domain combinations in the proteome. The pathogenesis of cancer can often be conceptualized as tumor cells rewiring and breaking mechanisms of known regulators of signaling networks to establish the various hallmarks of cancer like sustained proliferation or evasion of the immune system [44]. However, our DANSy analysis suggests that the rules of the proteome that guide feasible domain combinations are too fundamental to violate. Instead, cancer fusion genes may be further exploiting aspects of these rules to explore feasible domain combinations as seen with tyrosine kinase fusions. A major limitation of domain-abstracted networks though is it does not capture the specifics of a protein’s individual domain partnerships. More precisely, it can capture whether a domain combination already exists, but it does not capture whether the new partnerships, represented by that domain combination, introduce an important novelty into the network configuration. Hence, adaptations to n-gram networks that account for functional diversity on a partnership level, or complementary integration with protein-protein interaction networks will capture important biological insights, such as what will be needed to fully understand gene fusions and their role in disease.

## Materials and Methods

### Retrieving UniProt IDs for the human proteome and post-translational modification systems for additional species

For each species analyzed in this study, the canonical reference proteome was retrieved as a fasta file from the UniProt FTP server using the taxon ID and UniProt Reference Proteome IDs listed in Table S2. We retrieved the UniProt IDs for individual post-translational modification (PTM) systems by using the InterPro module of CoDIAC [26] for individual domains of each system using the InterPro IDs listed in Table S1. We repeated this process in the phosphorylation evolutionary analysis to get all possible candidates for each species, and then filtered UniProt IDs to those within their respective reference proteome.

### Fetching InterPro protein domain architectures and generating domain n-grams

To fetch protein domain architectures from InterPro, we utilized the UniProt module from CoDIAC [26] using the fetched UniProt IDs to create a reference file for all queried UniProt IDs. The resulting reference file contains all InterPro and UniProt domain architectures and reference sequences for each queried protein and was used for downstream analysis. All results were fetched using the InterPro version 108.0 and UniProt version 2026_01 builds of each database. The retrieved protein domain architectures were then separated into n-grams of the length of interest for each n-gram model. In the PTM analysis, the System Focused n-gram models, only n-grams that contained domains of interest were fetched from the complete protein domain architecture.

### Measuring n-gram model information gain and relative entropy

N-gram models rely on the Markov assumption where the probability of a specific n-gram depends on the conditional probability of the next domain, *d*_*n*_, given the preceding sequence of domains, *d*_*n−*(*N−*1):*n−*1_. This can be estimated for each n-gram using the maximum likelihood estimate (MLE):

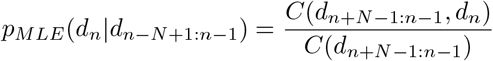

Where N represents the maximum length of n-grams being evaluated (i.e. N=2 for bigrams or N=5 for 5-grams). The counts of a specific n-gram (*d*_*n*+*N−*1:*n−*1_, *d*_*n*_) is represented by *C*(*d*_*n*+*N−*1:*n−*1_, *d*_*n*_). These probabilities are then used to calculate the entropy of an n-gram model (*H*_*n*_(*x*)) using Shannon’s entropy:

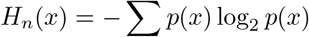

For individual domains the probabilities are given by the relative frequencies for each domain within the corpus of domain architectures. For longer n-grams, the entropy represents the sum of weighted probabilities which can be estimated by:

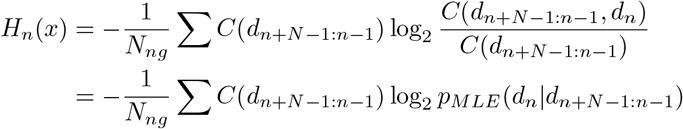

Where *N*_*ng*_ represents the total number of n-grams with length N. The entropy of an n-gram model is then used to determine the relative information gain *I*(*x*) from the unigram (only single domain frequency distributions) model (*H*_1_(*x*)):

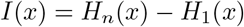

To compare how the distributions of n-grams change using different n-gram models we use the relative entropy also known as the Kullback-Leibler divergence defined as:

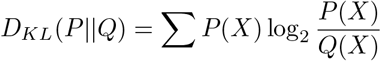

Where *P* (*X*) and *Q*(*X*) represent the probability distributions within both n-gram models and *Q*(*X*) is the baseline model that contains all n-grams within *P* (*X*).

### Calculating network centrality measurements

The betweenness and degree centrality measurements were calculated as implemented in the networkx python package [45]. Degree centrality is defined as the total fraction of all nodes connected to node *v*. The betweenness centrality measurement of node *v* is the sum of fractions of pairwise shortest paths that pass through node *v*, which is defined as:

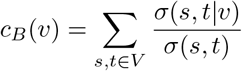

Where *V* is the set of nodes, *σ*(*s, t*) represents the number of shortest paths between nodes *s* and *t*, while *σ*(*s, t*|*v*) represents the number of shortest paths that go through node *v*.

### Predicting domain architectures for gene fusions

Genomic breakpoints for gene fusions were retrieved from the ChimerDB [34] for the TCGA cohort. Fusions were filtered to those predicted to remain in-frame or contain a portion of the coding region with breakpoints in either the 5’ or 3’ untranslated region and a part of the ChimerSeq+ dataset representing high confidence fusions. Each genomic breakpoint was converted to genomic coordinates of the current build of the human genome (hg38), and coordinates were deduplicated for any entries in ChimerDB for the same fusion gene in individual patient samples. For the CCLE dataset, we used publicly available cell line annotation from CCLE along with fusion calls and CRISPR knockout screening from the DepMap Public 26Q1 release (https://depmap.org/). Only high confidence, in-frame fusions were analyzed that had >0.1 fusion fragments per million total reads (FFPM) measures. Genomic breakpoints were then mapped to exon and the protein coding sequence positions. The base pair position was then translated to an amino acid position and used to determine which domains were donated towards the fusion gene (Fig. S10). Domains truncated by more than 5 amino acids (aa) for domains with an average size >50aa in size or 2aa for domains ≤ 50aa in length by the breakpoint location were excluded in the final predicted domain architecture.

### Domain n-gram enrichment analysis

For each collection of domain architectures, the enrichment of individual domain n-grams was calculated using a one-tailed hypergeometric test with the Benjamini-Hochberg to adjust p-values to control the false discovery rate. Domain n-grams were limited those with up to 3 domains in length. For our analysis of fusion genes, we further calculated the false positive rate (FPR) of the adjusted p-values using 100 trials of randomly chosen genes and breakpoints. For the TCGA cohort, we used a total of 15,000 gene pairs for each trial, while the CCLE cohort was compared to trials with 3000 pairs.

### Fusion gene dependency analysis

To calculate how fusion genes impact cell fitness, we retrieved the CRISPR knockout screening from the DepMap Public 26Q1 release (https://depmap.org/). For each fusion, we retrieved all cell lines from CCLE originating from the same lineage(s). Within each lineage, we calculated the mean difference in dependency score (MDDS) for each cell line by averaging the dependency score for each partner gene of a fusion gene. We then calculated the difference in the dependency scores between cells with the fusion and those without. Any fusion gene with an MDDS >0.5 was considered to be a potential dependent fusion gene similar to prior studies [46].

### Calculating Gini-Simpson Diversity Index for individual domains

To calculate the diversity index, we first annotated each domain architecture to include symbols for the N- and C-terminal ends to be extracted as part of individual domain n-grams (e.g. N-terminal|SH2). For each domain of interest *d*_*n*_ all 2-grams with the domain were retrieved and split into those that start or end with the domain of interest. For the n-grams starting with *d*_*n*_ were used to calculate the C-terminal diversity index, while those ending with *d*_*n*_ were used for the N-terminal diversity index. The diversity index was calculated using the Gini-Simpson Diversity Index as defined for small datasets:

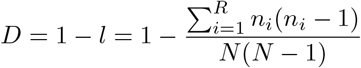

Where *R* is the collection of n+1 domain n-grams, *N* is the total count of the n-grams in the set, and *n*_*i*_ is the count for each individual domain n-gram.

### Species N-gram Similarity Index

For comparing the n-grams of the pTyr and pSer/Thr systems between individual species, the Jaccard similarity index, *J*, was calculated using:

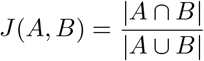

Where *A* and *B* represent the sets of domain n-grams found within individual species.

## Code and Data availability

The source code and installation guide for DANSy is provided at our Git repository: https://github.com/NaegleLab/DANSy. The specific applications and analysis performed with DANSy and other additional analyses are found at the Git repository: https://github.com/NaegleLab/DANSy_Applications.

## Acknowledgments

We would like to acknowledge the constructive and critical assessment of the presented work from the entire Naegle lab. The authors thank Bruce Mayer for helpful discussions about the phosphotyrosine system of *Monosiga brevicollis*. Research reported in this publication was supported by the National Institute of General Medical Sciences of the National Institutes of Health under Award Number R35GM138127. The content is solely the responsibility of the authors and does not necessarily represent the official views of the National Institutes of Health.

