## Supplementary Information for "Uncovering the domain language of protein function and protein networks using DANSy"

### Supplementary Information
#### Uncovering the domain language of protein functionality and cell phenotypes using DANSy.

Adrian A. Shimpi[1,2], Kristen M. Naegle[1,2,*]

**1 Department of Biomedical Engineering, University of Virginia, Charlottesville, VA, 22903**
**2 Department of Genome Sciences, University of Virginia, Charlottesville, VA, 22903**

##### This file includes:

##### Supplementary Text:

**Supplementary Note 1**: Expanded description of the entropic information of different n-gram models of the proteome.
**Supplementary Note 2**: Novel domain architectures of fusion genes impacts on DANSy networks.

##### Supplementary Figures:

**Supplementary Figure S1**: Extended n-gram network characterization of the human proteome.
**Supplementary Figure S2**: Isolates and Connected Components comparisons after collapsing redundant n-grams.
**Supplementary Figure S3**: Evaluation of different n-gram length models.
**Supplementary Figure S4**: pTyr domain distribution for individual species.
**Supplementary Figure S5**: pSer/Thr domain distribution for individual species.
**Supplementary Figure S6**: The complete set of the pTyr network across evolution.
**Supplementary Figure S7**: The complete set of the pSer/Thr network across evolution.
**Supplementary Figure S8**: Similarities of n-gram of the phosphorylation n-gram networks across species.
**Supplementary Figure S9**: Reversible PTM n-gram networks.
**Supplementary Figure S10**: Workflow for predicting gene fusion domain architectures.
**Supplementary Figure S11**: Network impacts of novel domain architectures.
**Supplementary Figure S12**: Relationship of breakpoints, unique fusion proteins, and unique predicted fusion domain architectures for TCGA cohorts.
**Supplementary Figure S13**: Breakpoint characterization for individual patients.
**Supplementary Figure S14**: DANSy analysis on the CCLE/DepMap dataset.
**Supplementary Figure S15**: Relationship of TCGA domain enrichments to network centrality metrics and protein counts.
**Supplementary Figure S16**: Expanded kinase fusion analysis in the CCLE dataset.

##### Supplementary Tables:

**Supplementary Table S1**: Reveresible PTM System Domain Categories and InterPro IDs.
**Supplementary Table S2**: Individual species names used to fetch UniProt IDs.

#### Supplementary Note 1: Expanded description of the entropic information of different n-gram models of the proteome.

When building the n-gram networks for DANSy, we aimed to balance the information being extracted (i.e. n-gram frequencies) with the network complexity to ensure sufficient representation of the proteome. As longer n-grams are extracted, we are gathering additional context for fewer n-grams in the proteome, which may have decreasingly less influence in the wider proteome and provide redundant information captured by the shorter n-grams. We addressed this concern partially by including the network collapsing step outlined in Figure 1C-D to reduce redundancy, but we wanted to determine the information content represented by different n-gram lengths. Here, we provide additional details on how we concluded that a 10-gram network model is sufficient for future DANSy-based analysis.

As outlined in the Methods, we determined the information encoded by different n-gram models by using the information gain $I(x)$ of each model relative to using only individual domain frequency distributions, and comparing that to the full network model. From these measurements, we found the simplest 2-gram model had an information gain of about 2 bits or almost 40% of the information in the full n-gram model. Information gains became more modest as longer n-grams were included in each model's corpus with the 5-, 10-, and 15-gram models having information gains of 3.77, 4.56, and 4.85 bits respectively or 71%, 85%, and 91% of the full n-gram model (Fig. S3A).

Given the information gain from the 2-, 5-, 10-, and 15-gram models (Fig.S3B), we asked if their respective n-gram networks recapitulated the network structure of the full n-gram model developed for Figure 1 by analyzing changes in the connected components. We found most (>75%) isolates from the original n-gram model were retained across each model. However, the 2-gram and 5-gram models also generated 771 and 28 additional isolates respectively (Fig.S3C), which suggests that the same set of proteins are spread across not only multiple nodes, but multiple connected components. We next analyzed the changes in non-isolate connected components, and found the 2-gram model only recapitulated half of the existing components in their entirety, and split 16 components into two or more additional components. Meanwhile, the 5-gram split 2 connected components and the 10- and 15-gram models only truncated the connected components by removing nodes associated with longer n-grams (Fig.S3D). The 15-gram model only truncated the largest connected component. Through this analysis, we ensure the properties that determine feasible domain combinations to endow protein function are represented and retained in models with 10-grams or longer.

Next, we wanted to determine whether the information content of each n-gram model diverges from the complete model to hinder our ability to capture meaningful connections of domain compatibility. In other words, does knowing the preceding 3-gram improve the probability of identifying the next domain in a 4-gram than only the preceding 2-gram, or are we increasing the complexity of n-grams being extracted without a concurrent increase in information. We quantify the collective difference in information content of each n-gram model through the relative entropy (or Kullback-Leibler divergence). Like the network changes, the 2-gram model exhibit the highest relative entropy (i.e. the largest difference), while the 5-, 10-, and 15-gram models had relative entropy values less than 0.4 bits with decreasing improvements for longer n-gram models (Fig. S3D)

Collectively, our results suggest that a 2-gram model insufficiently recapitulates the grammatical rules of the complete proteome. Despite large improvements in information, the changes of the network structure by the 5-gram model hinder its potential to approach the improvements of the longer 10- and 15-gram models. We observed similar trends in information and network structure between the 10- and 15-gram models with only minimal improvements for the 15-gram model, which do not offset the increased complexity. Thus, for the purposes of DANSy a 10-gram model is sufficient to recapitulate the grammatical rules seen within the full n-gram model of the complete proteome.

#### Supplementary Note 2: The network impacts of novel domain architectures from fusion genes.

For novel domain architectures, the addition of new nodes to the DANSy n-gram network can create two distinct changes to the network (Fig. S11A). The first occurs when a new node shortens existing paths between n-grams that have a common domain partner in the human network, or in other words are found within the same connected component of the network. An example of this is the BCR-ABL1 fusion gene found in chronic myelogenous leukemia. In the BCR-ABL1 fusion, BCR-oligomerization domain has no direct path to the protein kinase domain. Instead, to transit between the oligomerization domain and a protein kinase domain in the network, one must go through one of the n-gram nodes that includes either the DH, PH, or SH3 domains. By introducing the n-grams from the BCR-ABL1 fusion, we now introduce at least one n-gram (Bcr-oligomerization|DH|PH|SH3|SH2|Kinase) that contains both the oligomerization and kinase domain, thereby shortening the path between the domains. We call this subcategory “novel: shared” as impacts for the DANSy network. Alternatively, adding a node to the network can create a path between fully disconnected domain n-grams in the network. This is exemplified by the FGFR3-TACC3 fusion that results in a chimera protein that contains both a kinase domain and the TACC C-terminal domain. In the native proteome, the TACC C-terminal domain is an isolate and has no feasible path to the rest of the n-gram network including the kinase domain. By including the FGFR3-TACC3 domain n-grams, a path now exists between the two domains in the network that represents a new protein context that contains two previously obligate functions, which we term as “novel: obligate” for our DANSy n-gram network. Applying these subcategories to the novel domain architectures in the TCGA cohorts, we see that the “novel: shared” subcategory is the larger fraction for most cohorts. The cohorts with the fewest patients with fusions were exclusively the “novel: obligate” subcategory (Fig. S11B). Each of these subcategories creates new connections within the network that represent compatible biochemical or structural properties between the domains to create a feasible domain combination to create new protein functions and contexts not previously observed in the native proteome.

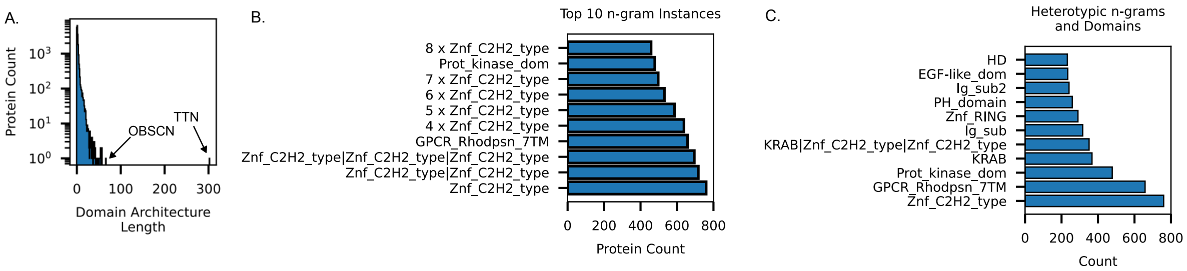


**Supplementary Figure S1. Extended n-gram network characterization of the human proteome.** A) Extended domain architecture length from Fig. 1E. B) Protein counts for the top domain n-grams or C) heterotypic n-grams or individual domains.

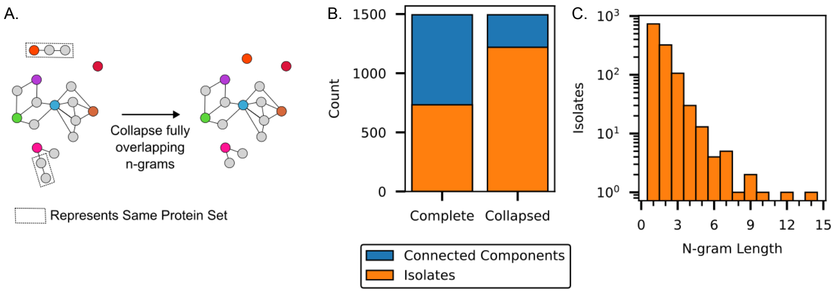


**Supplementary Figure S2. Isolates and Connected Components comparisons after collapsing redundant n-grams.** A) Schematic of collapsing n-grams that which fully represent the same set of proteins. B) Total count of isolates and connected components before and after collapsing n-grams representing the same protein families. C) Length of n-grams represented by the isolates after the collapse.

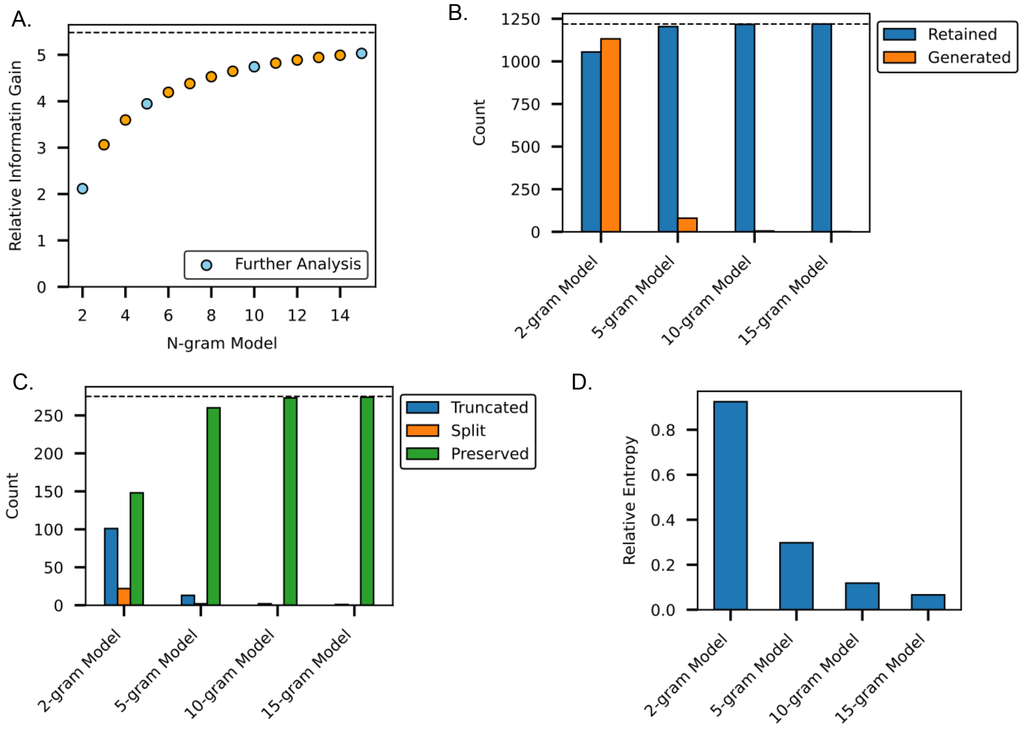


**Supplementary Figure S3. Comparison of different n-gram length models.** A) Information gain of different n-gram models relative to a unigram model. B) The number of isolates generated or retained in each individual n-gram model. C) The number of connected components that were either truncated, split, or fully preserved. D) The relative entropy of each n-gram model to the complete proteome model.

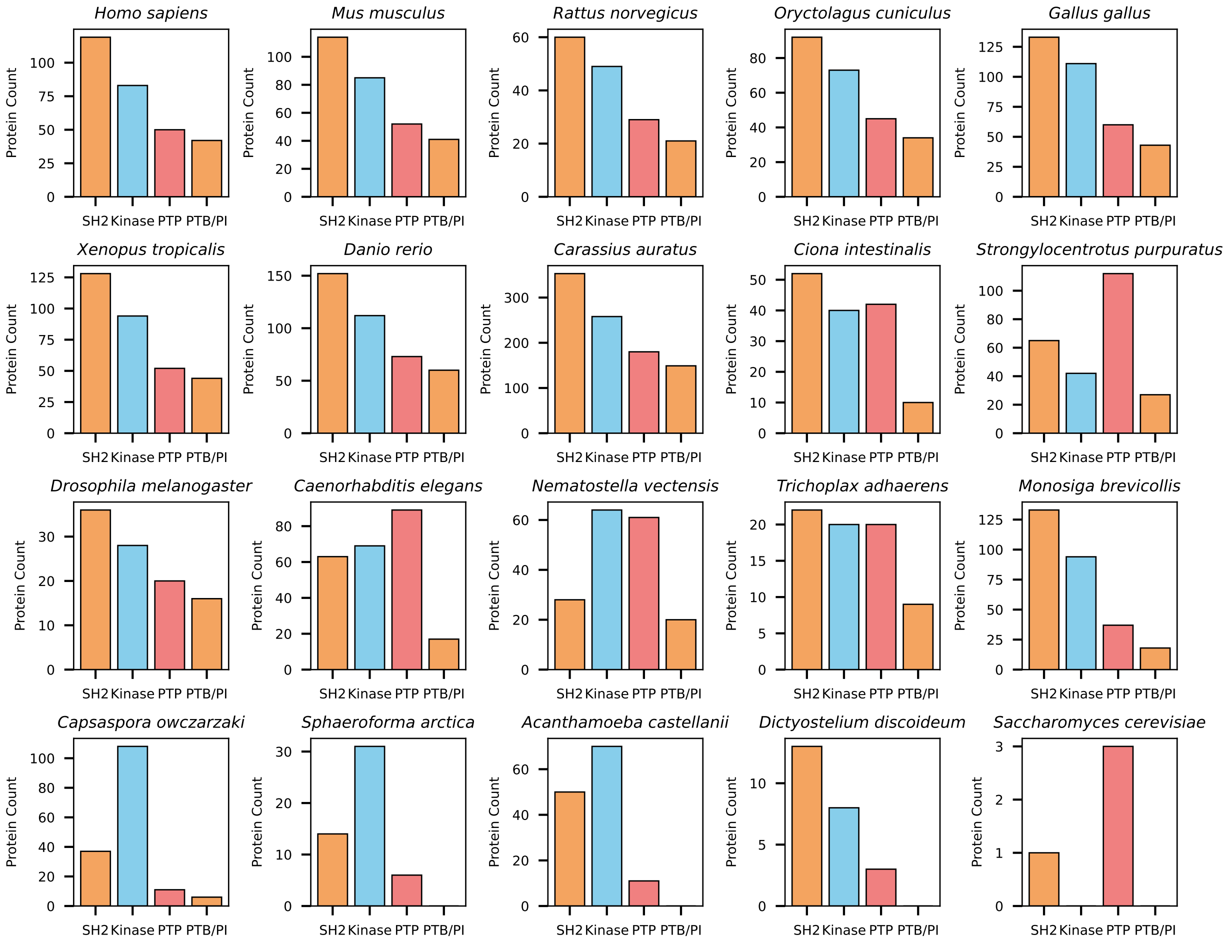


**Supplementary Figure S4. pTyr domain distribution for individual species.** Individual protein counts for each domain that constitutes the pTyr system.

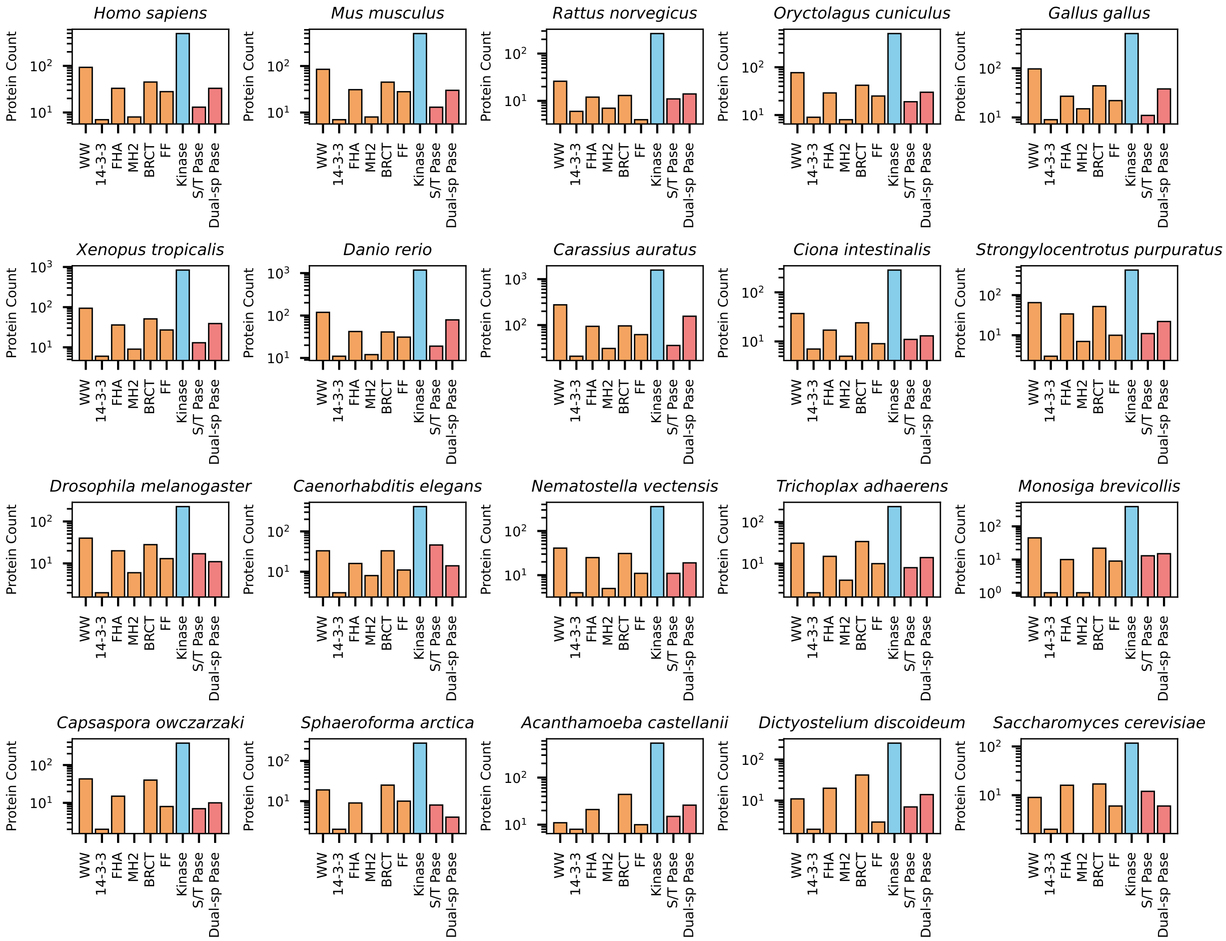


**Supplementary Figure S5. pSer/Thr domain distribution for individual species.** Individual protein counts for each domain that constitutes the pSer/Thr system.

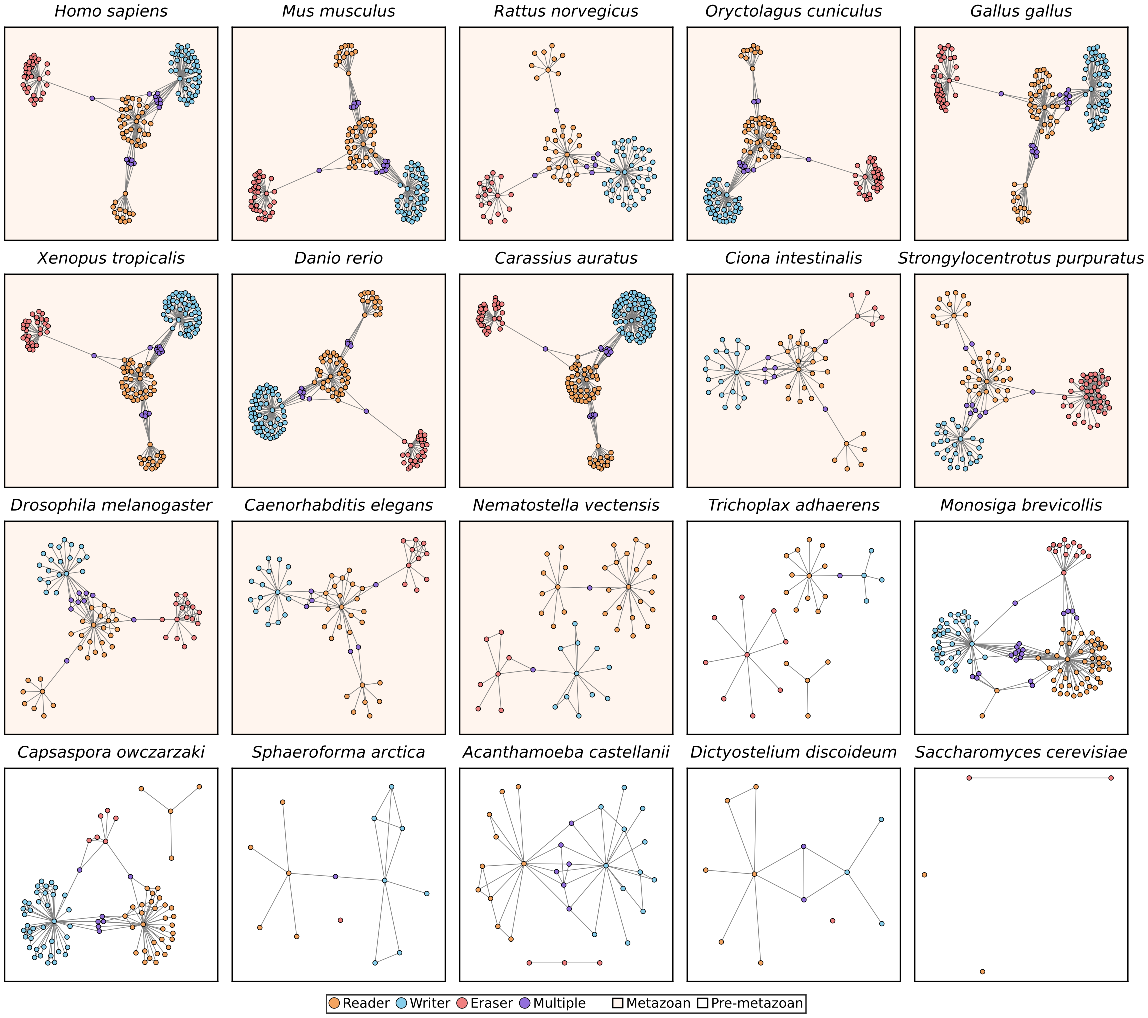


**Supplementary Figure S6. The complete set of the pTyr network across evolution.** The n-gram networks for the pTyr system for the complete list of species.

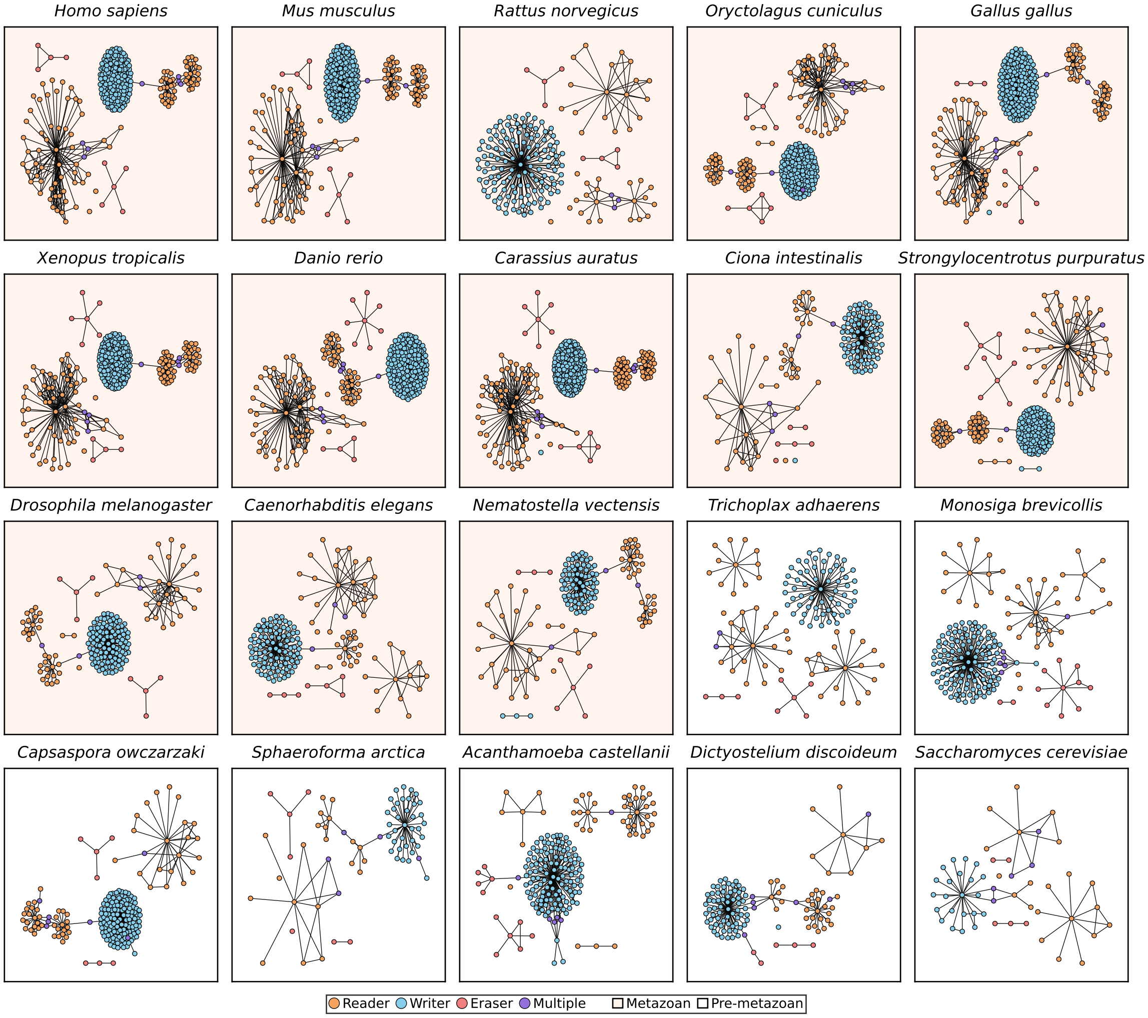


**Supplementary Figure S7. The complete set of the pSer/Thr network across evolution.** The n-gram networks for the pSer/Thr system for the complete list of species.

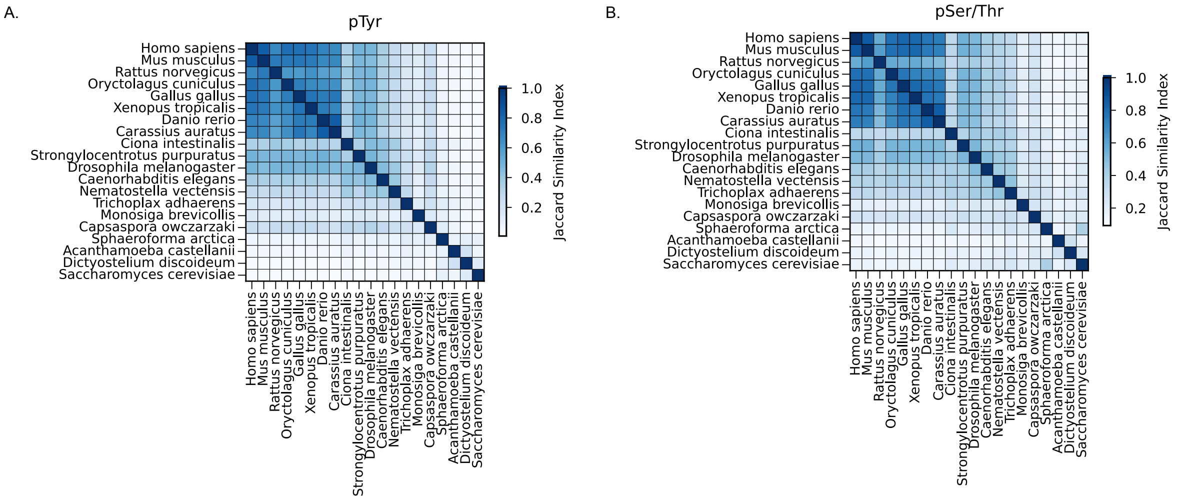


**Supplementary Figure S8. Similarities of n-gram of the phosphorylation n-gram networks across species.** The Jaccard similarity index of the n-grams extracted from each species for the A) phosphotyrosine or B) phosphoserine/threonine system.

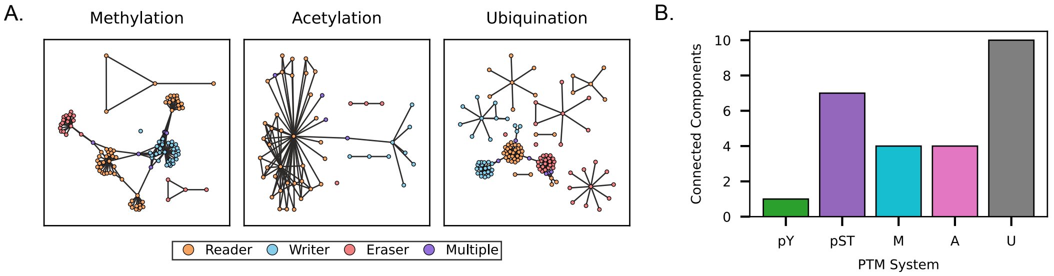


**Supplementary Figure S9. Reversible PTM n-gram networks.** A) The System Focused n-gram networks of proteins for additional reversible PTM systems including methylation, acetylation, and ubiquitination. B) The number of connected components for both the phosphorylation systems and additional reversible PTM systems which operate under a reader-writer-eraser paradigm. pTyr: Phosphotyrosine, pSer/Thr: Phosphoserine/threonine M: Methylation, A: Acetylation, Ub: Ubiquination

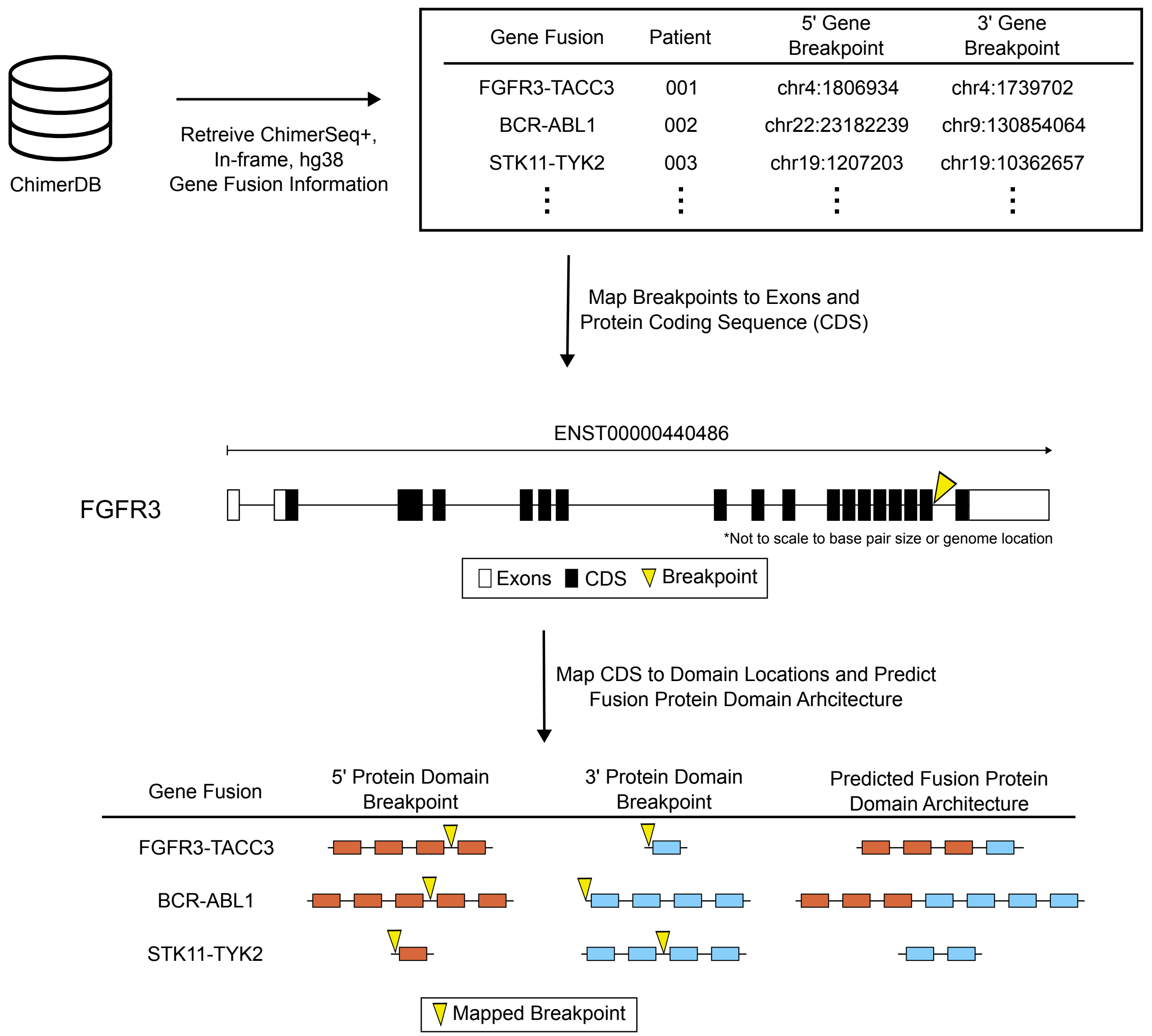


**Supplementary Figure S10. Workflow for predicting gene fusion domain architectures.** Individual fusion genes from ChimerDB were retreived and filtered to be in-frame, mapped to the hg38 genome build, and are high confidence fusions as included within ChimerSeq+. Genomic breakpoints were mapped to the exon and protein coding sequence positions, translated to the amino acid position, and used to determine domains donated by parent genes.

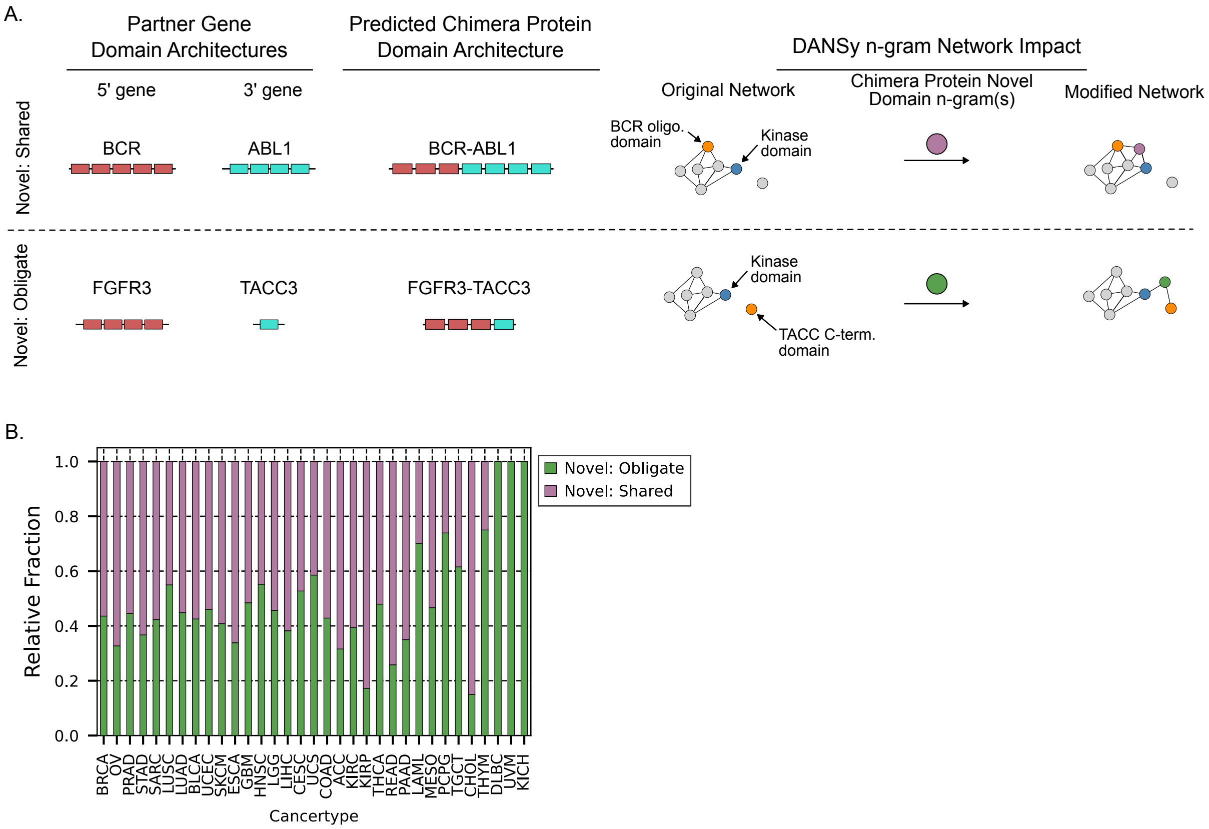


**Supplementary Figure S11. Network impacts of novel domain architectures.** A) Schematic of potential domain n-gram network impacts created by adding nodes representing n-grams from novel domain architectures from fusion genes. B) The relative fraction each novel domain subcategory in each TCGA cohort.

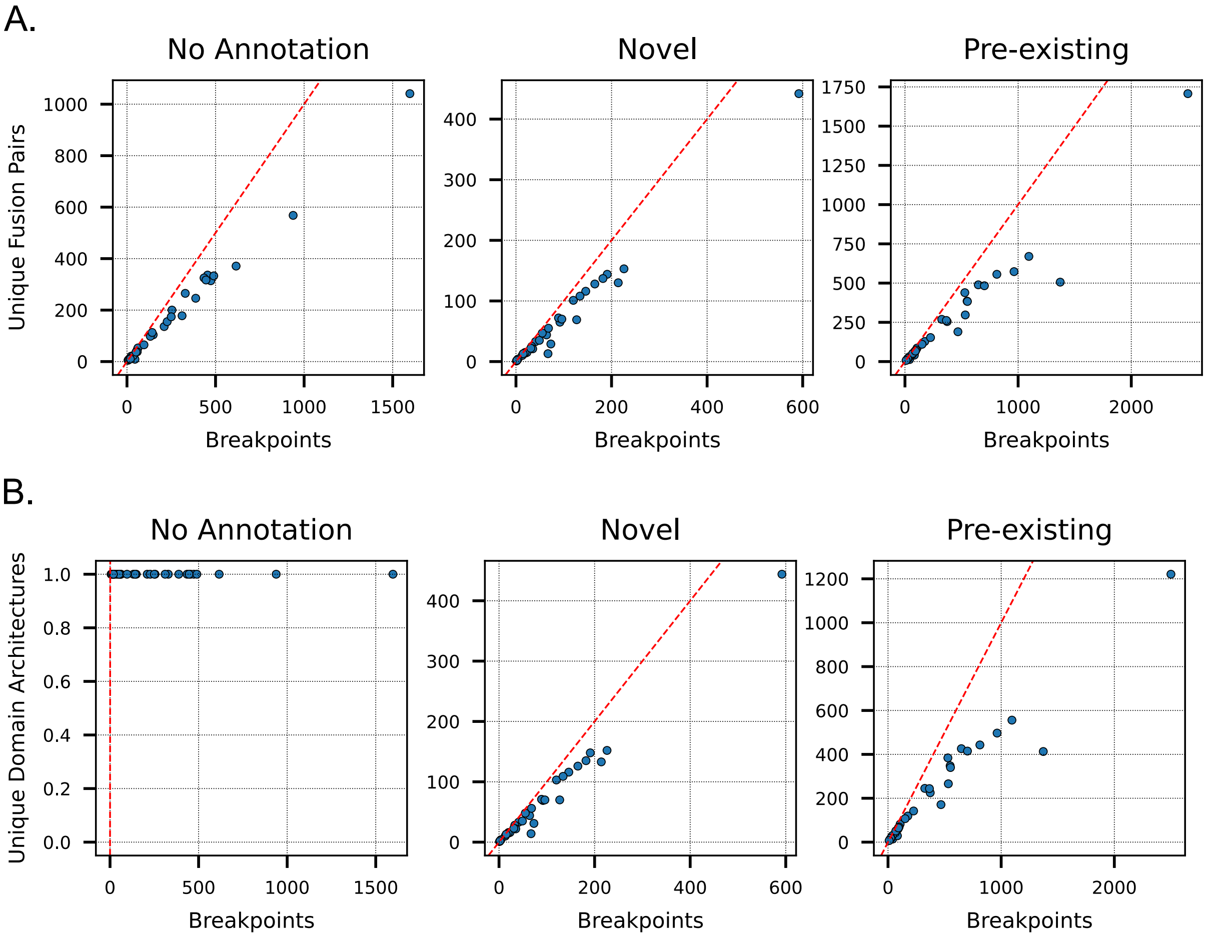


**Supplementary Figure S12. Relationship of breakpoints, unique fusion proteins, and unique predicted fusion domain architectures for TCGA cohorts.** For each TCGA cohort, the relative relationship of A) number of breakpoints to unique fusion partner gene pairs, B) number of breakpoints to unique predicted domain architectures, or C) number of unique domain architectures to unique fusion partner gene pairs for each DANSy category.

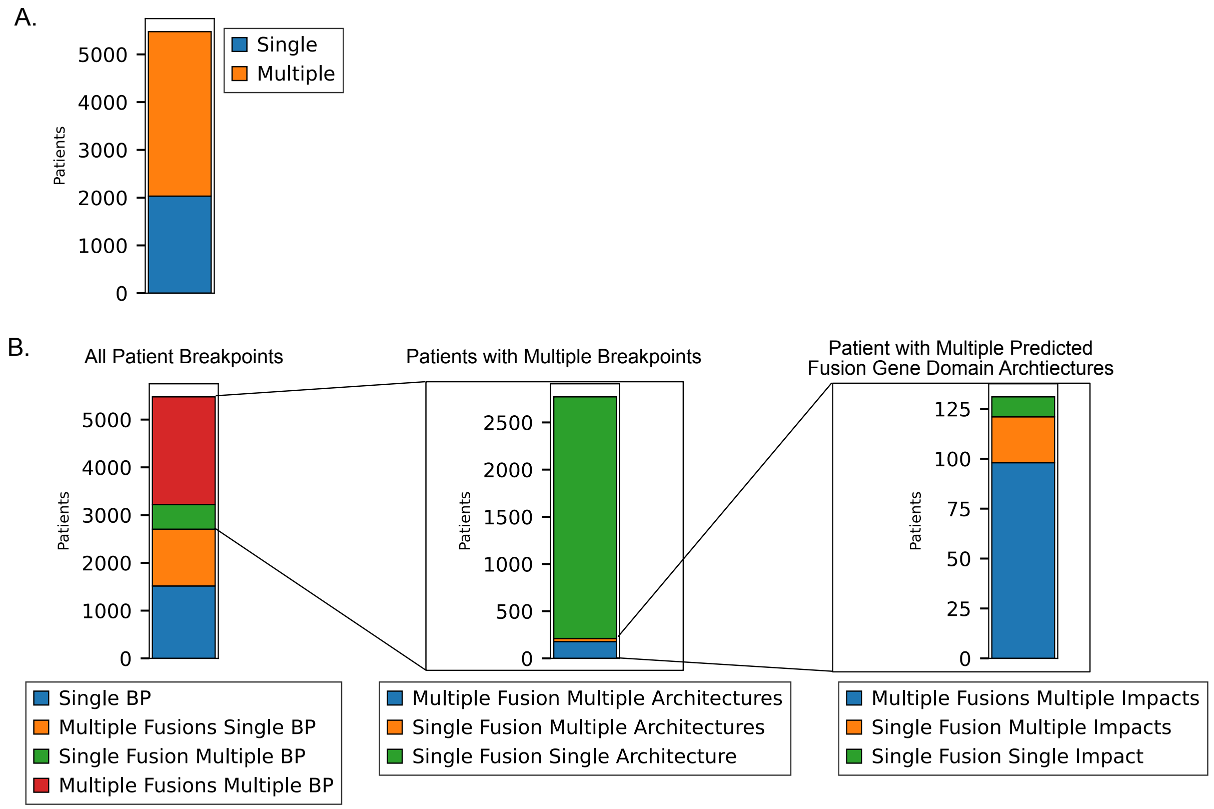


**Supplementary Figure S13. Breakpoint characterization for individual patients.** A) The distribution of patients with either single or multiple fusion gene breakpoints identified across the TCGA cohort. B) The distribution of patients and if they have a single or multiple fusion genes with single or multiple breakpoints. C) The distribution of patients with multiple breakpoints and their relationship to generating single or multiple domain architectures. D) The distribution of patients with multiple domain architectures for a single fusion gene pair and whether the domain architecture changed the DANSy category.

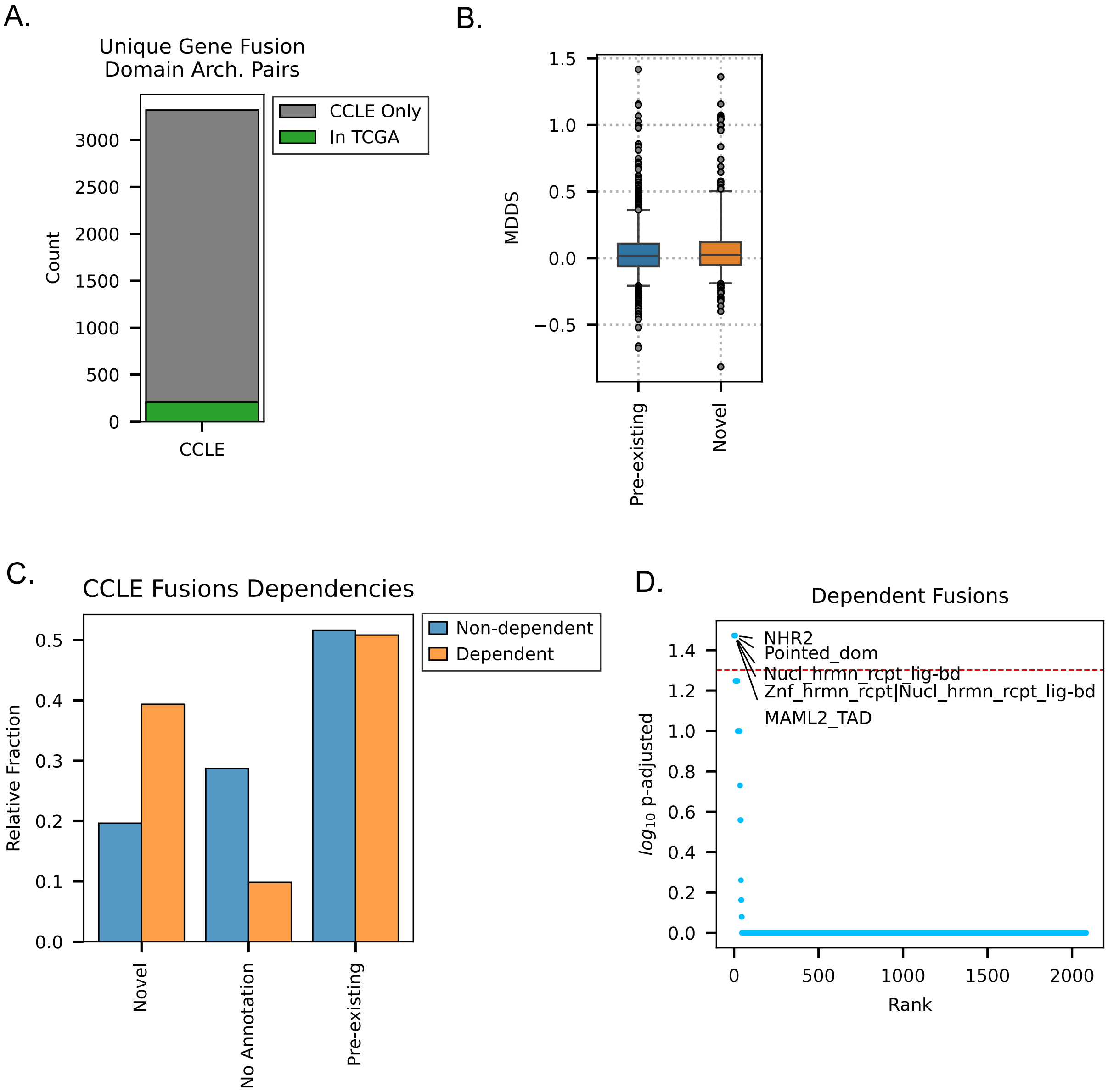


**Supplementary Figure S14. DANSy analysis on the CCLE/DepMap dataset.** A) The number of unique gene fusion-domain architecture pairs in the CCLE and their overlap with TCGA. B) The distribution of the mean differential dependency scores (MDDS) for fusion genes that either created pre-existing or novel domain architectures. C) The distribution of DANSy categories for fusion genes that present a possible dependency. D) The domain n-grams enriched in cell lines from DepMap that exhibit a potential dependence for the fusion gene.

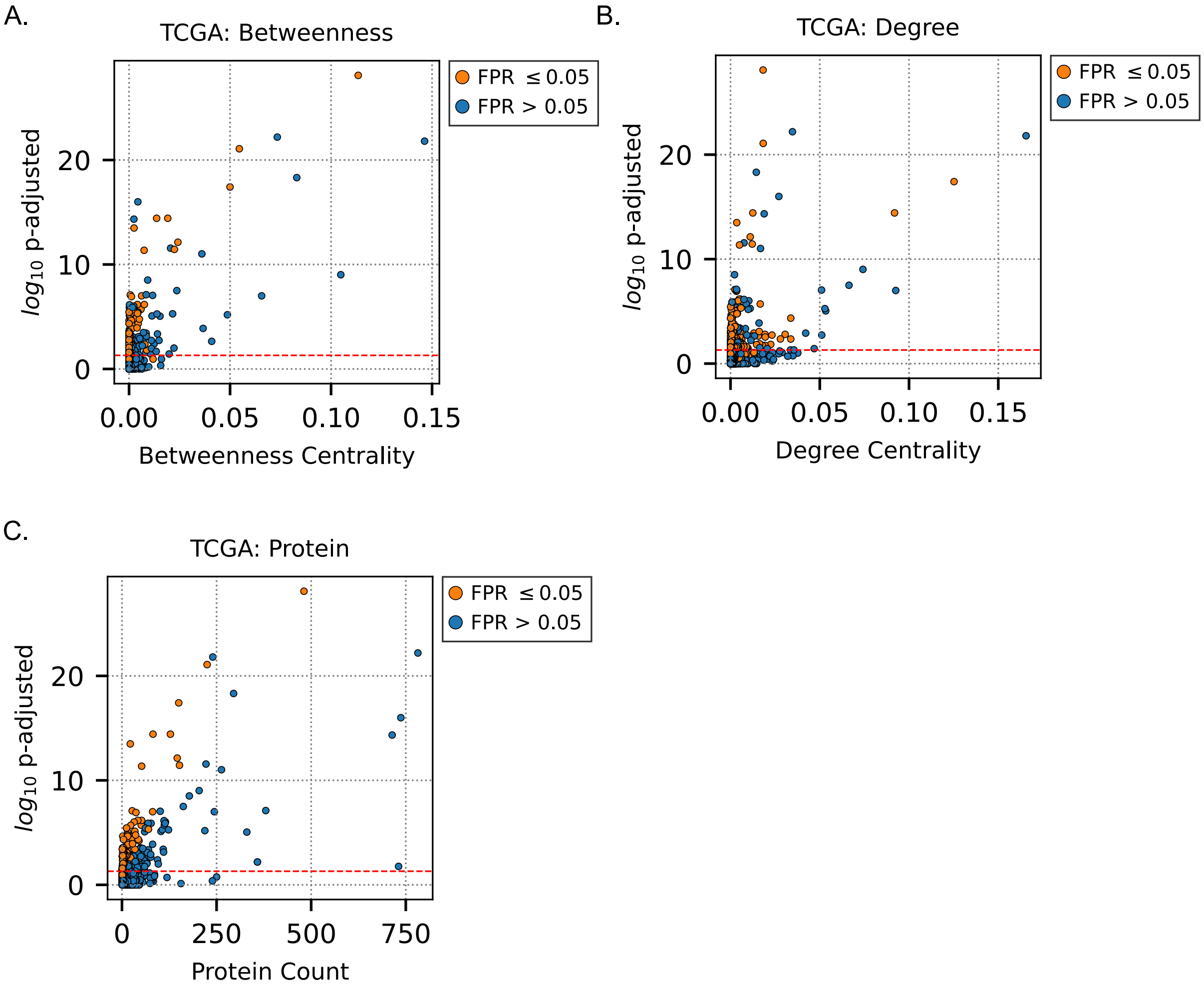


**Supplementary Figure S15. Relationship of TCGA domain enrichments to network centrality metrics and protein counts.** The correlation between the domain n-gram enrichment values relative to A) betweenness centrality, B) degree centrality, or C) the number of proteins with the domain. The distribution of each value is displayed as histogram on the opposite axis. For each plot the dashed red line indicates an adjusted p-value of 0.05.

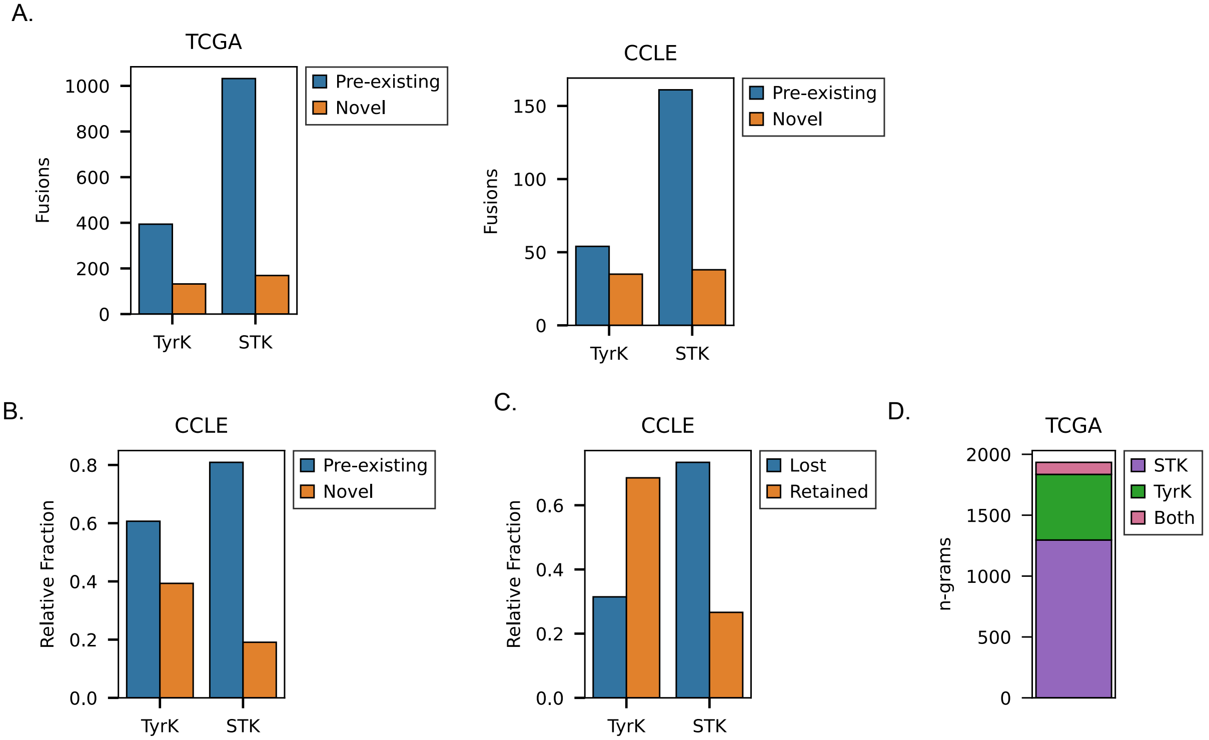


**Supplementary Figure S16. Expanded kinase fusion analysis in the CCLE dataset.** A) The number of either tyrosine kinase (TK) or serine/threonine kinase (STK) fusions within the TCGA (left) or CCLE (right) datasets and whether they create novel or pre-existing domain architectures. B) The relative fraction of TK or STK fusions creating novel domain architectures in the CCLE dataset. C) The relative fraction of TK or STK fusions in CCLE that retain the kinase domain in their predicted domain architecture.

**Supplementary Table S1. Reveresible PTM System Domain Categories and InterPro IDs.**

| PTM System | Reader | Writer | Eraser |
|---|---|---|---|
| **pTyr** | SH2 (IPR000980)<br>PTB/PI (IPR006020) | Tyr-Kinase (IPR020635) | PTP_Catalytic (IPR000242) |
| **pSer/Thr** | WW (IPR001202)<br>14-3-3 (IPR023410)<br>FHA (IPR000253)<br>MH2 (IPR001132)<br>BRCT (IPR001357)<br>FF (IPR002713) | Ser/Thr Kinase (IPR001245) | Ser/Thr Phosphatase (IPR006186)<br>PPM-type Phosphatase (IPR001932)<br>FCP1-Homology (IPR004274)<br>Dual-specificity Phosphatase (IPR000340) |
| **Acetylation** | Bromodomain (IPR001487) | MYST (IPR002717)<br>CBP (IPR031162)<br>TAF1 (IPR022591)<br>GNAT (IPR000182) | HDAC (IPR023801)<br>Sirtuin Catalytic Core (IPR026590) |
| **Methylation** | TUDOR (IPR002999)<br>Agenet (IPR008395)<br>PWWP (IPR000313)<br>Chromodomain (IPR023780) | SET (IPR001214)<br>DOT (IPR025789) | Amine oxidase (IPR002937)<br>JmjC (IPR003347) |
| **Ubiquination** | UBA (IPR015940)<br>CC2-LZ (IPR032419)<br>GAT (IPR004152)<br>CUE (IPR003892)<br>Znf-UBZ (IPR041641)<br>Znf-UBP (IPR001607) | HECT (IPR000569)<br>Thif-type NAD/FAD binding fold (IPR000594)<br>UBQ conjugating E2 (IPR000608) | OTU (IPR003323)<br>USP (IPR028889)<br>JAMM/MPN+ (IPR000555)<br>Josephin (IPR006155) |

**Supplementary Table S2. Individual species names used to fetch UniProt IDs.**

| **Species** | **Strain** | Proteome ID | Taxon ID |
|---|---|---|---|
| *Homo sapiens* | | UP000005640 | 9606 |
| *Mus musculus* | | UP000000589 | 10090 |
| *Rattus norvegicus* | | UP000002494 | 10116 |
| *Oryctolagus cuniculus* | | UP000001811 | 9986 |
| *Gallus gallus* | | UP000000539 | 9031 |
| *Xenopus tropicalis* | | UP000008143 | 8364 |
| *Danio rerio* | | UP000000437 | 7955 |
| *Carassius auratus* | | UP000515129 | 7957 |
| *Ciona intestinalis* | | UP000008144 | 7719 |
| *Strongylocentrotus purpuratus* | | UP000007110 | 7668 |
| *Drosophila melanogaster* | | UP000000803 | 7227 |
| *Caenorhabditis elegans* | | UP000001940 | 6239 |
| *Nematostella vectensis* | | UP000001593 | 45351 |
| *Trichoplax adhaerens* | | UP000009022 | 10228 |
| *Monosiga brevicollis* | | UP000001357 | 81824 |
| *Capsaspora owczarzaki* | ATCC 30864 | UP000008743 | 595528 |
| *Sphaeroforma arctica* | JP610 | UP000054560 | 667725 |
| *Acanthamoeba castellanii* | ATCC 30010 / Neff | UP000011083 | 1257118 |
| *Dictyostelium discoideum* | | UP000002195 | 44689 |
| *Saccharomyces cerevisiae* | ATCC 204508 / S288c | UP000002311 | 559292 |
